# Prion Seeding Activity in DNA Extractions: Implications for Laboratory Biosafety

**DOI:** 10.1101/2025.09.15.676309

**Authors:** Sarah C. Gresch, Tamara Morrill, Maddy Ellis-Cramer, Maria Arifin, Lexi E. Frank, Jason C. Bartz, Marc D. Schwabenlander, Tiffany M. Wolf, Gordon B. Mitchell, Jiewen Guan, Peter A. Larsen

## Abstract

Infectious prions (PrP^Sc^) are largely resistant to proteolytic digestion, including proteinase K digestion. While nucleic acid extracts are generally considered non-infectious, we investigated whether standard DNA purification methods can co-purify PrP^Sc^, posing an unrecognized biosafety risk. Two laboratories, the University of Minnesota Center for Prion Research and Outreach (MNPRO) and the Canadian Food Inspection Agency (CFIA), independently tested filter-based and magnetic bead-based DNA extraction kits using tissues from chronic wasting disease (CWD)-positive and -negative white-tailed deer (WTD; Odocoileus virginianus), as well as prion-infected and control Syrian hamster (Mesocricetus auratus) brains. CFIA used two filter-based kits (one automated and one manual), while MNPRO tested two manual kits (one filter-based and one magnetic bead-based). PrP^Sc^ seeding activity was measured in extracted DNA and source tissues using real-time quaking-induced conversion (RT-QuIC). MNPRO found substantial to almost perfect agreement (kappa (κ) = 0.789 - 0.816) between RT-QuIC seeding activity of DNA eluates from both extraction methods and that of the source WTD tissue homogenate. CFIA optimized RT-QuIC to a 30-hour runtime, achieving 74% sensitivity and 94% specificity in 88 archived WTD DNA samples. Both laboratories concluded that commercial DNA extraction kits do not eliminate PrP^Sc^, enabling its carry-over into DNA eluates. Until infectivity is resolved by animal bioassay, DNA from PrP^Sc^-positive tissues should be managed under biosafety protocols appropriate for the originating prion disease, with appropriate decontamination and containment procedures.

## Introduction

Infectious prions (PrP^Sc^) are misfolded proteins that can induce abnormal folding of normal cellular prion proteins (PrP^C^), leading to transmissible spongiform encephalopathies (TSEs) in both animals and humans. A defining characteristic of prions is their remarkable resistance to conventional decontamination methods, including heat, radiation, chemical disinfectants, and proteolytic enzymes, including proteinase K (PK) [1–4]. This resilience poses challenges in laboratory environments, necessitating stringent biosecurity protocols to prevent inadvertent exposure, cross-contamination, and environmental release of prion-contaminated material. Recent cases of variant Creutzfeldt–Jakob disease (vCJD) linked to accidental laboratory exposure to bovine spongiform encephalopathy (BSE) prions underscore the ongoing need for vigilance and enhanced containment in research and diagnostic settings [5, 6].

Many PrP^Sc^ diagnostic techniques, including Western Blotting, enzyme-linked immunosorbent assay (ELISA), protein misfolding cyclic amplification (PMCA) assay, mass spectrometry, and immunoassay, use PK digestion to distinguish PrP^C^ from PrP^Sc^ [7–10]. PK digestion methods are also commonly used during nucleic acid purification protocols. While launching new genetic and diagnostic analyses focused on chronic wasting disease (CWD), our teams independently questioned whether PrP^Sc^ could persist through standard DNA extraction protocols and remain intact within DNA eluates.

Prion proteins are known to interact with nucleic acids in vitro, with both PrP^C^ and PrP^Sc^ binding to DNA and RNA via electrostatic interactions [11, 12]. These associations can promote structural rearrangements in PrP and have been implicated in prion aggregation, conversion, and strain propagation [13, 14]. PrP^Sc^–nucleic acid complexes have also been described by Zou et al. and Nandi & Nicole, who demonstrated specific binding to DNA in biochemical assays [15, 16]. However, these studies did not evaluate whether nucleic acid-associated PrP^Sc^ retains seeding ability, leaving open questions about the biological relevance of these interactions.

To our knowledge, no prior study has used a prion amplification assay to evaluate whether DNA eluates from prion-infected tissues harbor PrP^Sc^ capable of seeding activity. This has significant implications: from a viral or bacterial standpoint, nucleic acid extracts are typically regarded as non-infectious [17]. However, if PrP^Sc^ retains seeding activity in these eluates, then DNA from TSE-positive samples may warrant biosafety controls consistent with the prion disease of origin. This is especially relevant given the growing efforts to better understand genetic risk factors associated with both human and animal TSEs. Many studies on CJD [18, 19], BSE [20, 21], scrapie [22], and CWD [23–25] use extracted DNA for a variety of downstream research projects (e.g., PRNP genotyping, methylation studies, whole genome sequencing, and SNP arrays), presumably conducted across multiple research laboratories (e.g., external sequencing or genotyping core labs). Given the potential for PrP^Sc^ to persist in nucleic acid extracts, it is critical to evaluate nucleic acid samples from TSE-positive tissues to determine the presence or absence of PrP^Sc^. Such information would directly inform appropriate containment measures to mitigate the risk of inadvertent PrP^Sc^ exposures as well as environmental contamination.

In this study, we investigated whether commonly used DNA extraction methods (which include PK treatment) could inadvertently retain PrP^Sc^ in DNA eluates from CWD-positive white-tailed deer (WTD; *Odocoileus virginianus*) and hamster brains infected with synthetic prions [26]. Our investigations included both filter- and magnetic bead-based commercial DNA extraction kits. To test for the presence of PrP^Sc^ in DNA, we used real-time quaking-induced conversion (RT-QuIC) assays with appropriate negative controls, following standard CWD-testing procedures. Experiments were independently replicated in two laboratories, including blinded assessments, to ensure rigor and reproducibility across DNA extraction methods and tissue sources.

## Materials and Methods

Two independent research teams, one at the University of Minnesota Center for Prion Research and Outreach (MNPRO) and the other at the Canadian Food Inspection Agency (CFIA), conducted separate experiments to investigate the potential co-extraction of PrP^Sc^ with DNA. Both teams employed two DNA extraction protocols across a variety of tissue types and assessed the seeding activity of DNA extracts using RT-QuIC. Methods for each team are presented separately.

### MNPRO

#### Experimental design

MNPRO extracted DNA from 53 samples representing 21 tissue types collected from nine CWD-positive deer (Table A2). For comparison, 23 samples representing six tissue types were extracted from 11 CWD-negative deer. Each sample underwent two independent extractions (details below). The tissue homogenate and each DNA eluate were tested for seeding activity by RT-QuIC.

#### Sample information

WTD tissue samples were obtained from the MN DNR and MN hunters between 2019 and 2020, and have been stored at −80°C in MNPRO’s biorepository since then. Samples used for DNA extraction consisted of both CWD-positive and -negative tissues. Samples, with associated metadata, are listed in Tables A1-A3. A prescapular lymph node homogenate from a CWD-positive WTD was used as the positive control throughout this study. The negative control consisted of parotid lymph node homogenate from a CWD-negative WTD.

Hamster brain homogenates were provided through ongoing collaborations with the laboratory of Dr. Jason Bartz. Eight homogenates were derived from hamsters experimentally inoculated with synthetic PrP^Sc^, and nine were inoculated with sham control [26]. The lab technicians working on this study were blinded to the presence or absence of PrP^Sc^ in individual hamster brain samples until after testing and analyses were complete. All samples were handled assuming they were PrP^Sc^-positive.

#### Determination of CWD status

The CWD status of each WTD included in MNPRO experiments was previously established by ELISA screening with immunohistochemistry (IHC) confirmation of the retropharyngeal lymph node (RPLN; Tables A1 and A2). ELISA and IHC testing were performed for regulatory or management purposes by an independent laboratory (Colorado State University Veterinary Diagnostic Laboratories, National Veterinary Services Laboratories, University of Minnesota Veterinary Diagnostic Laboratory, or Wisconsin Veterinary Diagnostic Laboratory).

#### DNA extraction and quantification

DNA extraction methods were performed using the filter-based DNeasy Blood and Tissue Kit (Qiagen, cat. # 69504) or the magnetic bead-based MagAttract HMW DNA Kit (Qiagen, cat. # 67563). Extraction methods largely followed manufacturers’ protocols, except for subsample volume, where we used 40-50 mg of tissue instead of the recommended 25 mg of tissue. Every extraction included PK and RNase steps, following the manufacturer’s recommendations, to ensure the removal of RNA and any proteins sensitive to enzymatic degradation. Nuclease-free water was used for the final DNA elution for both extraction kits.

DNA was eluted twice per sample, at 100 uL each, and these elutions were combined so that each sample produced 200 uL of eluent. Qubit double-stranded DNA kits (Invitrogen, cat. # Q32850 and Q32851) were used to quantify the DNA concentration in the eluates. DNA concentrations for individual samples are shown in Tables A1-A3.

#### Endpoint Titration Experiments

A subset of ten tissue samples (nine CWD-positive and one CWD-negative) was analyzed by endpoint titration to assess relative prion titer levels. Tissue homogenates and the DNA extracted from them were included in these experiments. Homogenates underwent RT-QuIC testing in serial dilutions ranging from 10⁻³ to at least 10⁻⁷, and 10^-8^ when needed, while extracted DNA was tested in serial dilutions from 10⁻¹ to 10⁻⁵. For these 10 samples, every dilution level of each homogenate and DNA eluate was tested in four technical replicates. The endpoint was defined as the final dilution level that showed statistically significant seeding activity (as described below), with dilutions tested at one to four levels beyond that, to ensure a true endpoint had been achieved.

#### RT-QuIC

MNPRO’s RT-QuIC assay parameters used herein are described by Schwabenlander et al. [27] with slight modifications consisting of shaking at 900 rpm, and shake/rest timing (60 seconds shaking, 60 seconds rest), repeated for 65 cycles (48 hours). All RT-QuIC reactions were performed on 96-well plates with CWD-positive and negative controls (n = 6 per control). A 100 mg (+/- 10 mg) subsample of tissue was added to 900 uL PBS and homogenized in a 2.0 mL tube containing 1.5 mm zirconium beads (cat. # D1032-15; Benchmark Scientific), using a BeadBlaster (Benchmark Scientific). All tissues were tested for seeding activity using RT-QuIC in a dilution buffer. BMG Omega Fluostar plate readers generated real-time fluorescence data. All RT-QuIC experiments were run at 42°C for 48 hours. DNA eluates were tested in four technical replicates by RT-QuIC, 1:1, using a dilution buffer (0.01% N2 supplement in a solution of 0.1% sodium dodecyl sulfate (SDS) in PBS). Aside from the endpoint titration experiments explained previously, tissues were tested at 10^-3^, except for muscle, which was tested at 10^-1^.

#### Data analysis

RT-QuIC data were analyzed using three assay metrics to determine whether samples were positive or negative. Ordinary one-way ANOVA with Fisher’s LSD test was conducted in GraphPad Prism (v10.2.3; graphpad.com) to compare the maxpoint ratio (MPR) and maximum slope (MS) against negative plate controls. A sample was classified as positive if at least 50% of its rate of amyloid formation (RAF) results were non-zero and at least one of the other metrics showed statistical significance (α = 0.05). RAF was calculated as the reciprocal of the time required for fluorescence to reach twice the background fluorescence. Background fluorescence was determined by averaging the fluorescence readings from cycles two and three to account for variability among wells. If a well did not reach the threshold, its RAF value was set to zero. MPR was defined as the maximum relative fluorescence units (RFU) divided by background RFU. MS was calculated as the greatest difference in RFU between the current time point and six cycles (∼4.5 hours) later, divided by time.

Agreement between categorical outcomes (CWD status, tissue homogenate seeding activity, and DNA eluate seeding activity) was evaluated using Cohen’s kappa (κ) with standard errors and 95% confidence intervals.

### CFIA

#### Experimental design

CFIA analyzed archived DNA from either obex or RPLN of 88 WTD (Table B2). DNA had been previously extracted using one of two methods (described below), so each tissue sample was represented by a single DNA eluate tested for RT-QuIC seeding activity.

CFIA first optimized the run-time of their RT-QuIC assay (results below) before testing DNA eluates.

#### Sample information

Obex and RPLN tissue specimens from WTD were received for CWD diagnostic testing and PRNP genotyping at the CWD Reference Laboratory, Canadian Food Inspection Agency. A random subset of specimens collected from provincial wildlife surveillance programs between January 2020 and February 2024 was selected for this study to determine if the DNA extracted from the tissue homogenates contain any prion seeding activity using RT-QuIC assays. Metadata associated with this subset is listed in Table B2.

#### Determination of CWD status

To determine the CWD status, tissue specimens were trimmed and homogenized to form a 10∼15% w/v homogenate, as described by Yilmaz et al. [28]. These homogenates underwent initial screening by ELISA using the TeSeE SAP Combi kit (Bio-rad, Hercules, CA, USA), followed by confirmatory testing by IHC [29]. Based on the CWD status, selected obex and RPLN tissue homogenates were used to generate control samples for optimizing RT-QuIC assays. CWD-positive homogenates were serially diluted 10-fold in CWD-negative homogenates to produce a concentration gradient ranging from 10^-5^ to 1.0^-1^ (v/v). As such, all control samples were 10∼15% w/v tissue homogenates containing various concentrations of PrP^Sc^.

#### DNA extraction and quantification

Obex and RPLN tissue homogenates and the above control samples, 80 ul each, underwent DNA extraction. DNA was extracted either using the QIAamp 96 DNA QIAcube HT Kit (Qiagen, cat. # 51331) on the QIAcube HT automated platform (Qiagen) or via manual extraction with the DNeasy Blood & Tissue Kit (Qiagen, cat. # 69504) according to the manufacturer’s instructions. Detailed information on the extraction method used for each homogenate was provided in Tables S1 and S2. For each sample, DNA was eluted in 150 µL AE buffer (Qiagen) and quantified using the Quant-iT dsDNA high sensitivity Assay Kits (Invitrogen, cat. # Q33120).

#### RT-QuIC

The RT-QuIC assay protocol and parameters used in this study were based on previously published methods [28], with slight modifications. In brief, each RT-QuIC reaction consisted of 5 µL of testing sample and 95 µL of master mix solution. To prepare the sample, tissue homogenates were diluted to 10^-4^ (w/v) in 0.05% SDS, and DNA eluate was added with 1% SDS in a ratio of 19:1 so that the testing DNA sample contained 0.05% SDS. The master mix solution contained 50 mM sodium phosphate buffer at pH 7.4, 300 mM NaCl, 1 mM EDTA, 10 μM Thioflavin T (ThT), and 0.1 mg/mL recPrP^C^ (Priogen Corp, Minnesota, United States). RT-QuIC reactions were run using BMG FluoStar® plate readers (BMG Labtech, Germany). At least eight replicates were tested for each control tissue homogenate, and 16 replicates for each corresponding DNA sample. DNA samples extracted from the archived tissue homogenates were tested in four or eight replicates, depending on the testing consistency as described below.

Assays were performed at 42°C for 65 hours to determine the optimal assay duration for DNA samples or using the optimized assay durations to detect prion seeding activity in DNA samples. Each cycle lasted approximately 17 minutes, with seven repeats of a one-minute shake at 700 rpm (double orbital) and a one-minute rest, followed by a one-minute reading. ThT fluorescence measurements were taken every cycle at a gain of 1200, excitation of 450 nm, and emission of 480 nm. Following the completion of the assay, the data were exported from the Mars data analysis software (BMG Labtech) and processed in Microsoft Excel and RStudio.

#### Data analysis

The RT-QuIC data were analyzed using a cycle threshold (CT) method as described by Yilmaz et al. [28]. Cycle threshold was defined as the time when the ThT signal of a reaction crossed the threshold, which was calculated using the average fourth cycle reading of all the reactions in a 96-well plate in RFU plus ten standard deviations. A CT of 65 hours was assigned for reactions where the ThT signal did not cross the threshold within the 65-hour assay. An RT-QuIC reaction was considered positive if the CT value was lower than or equal to the optimal assay duration determined for a specific sample type. For example, 33 h was used for obex and 30 h for RPLN [28].

#### Determination of optimal assay duration for DNA samples

To determine the optimal RT-QuIC assay duration for DNA samples, the approaches described by Yilmaz et al. [28] were followed and modified. In brief, CT values were generated with the control tissue homogenates containing CWD+ and CWD-tissue homogenates and their mixtures in a CWD+ to CWD-ratio ranging from 10^-5^ to 10^-1^ (v/v). Each control homogenate was classified as CWD+ if at least 4 out of 8 replicates in RT-QuIC showed a positive CT value; otherwise, it was classified as CWD-.

To determine the optimal assay duration time for DNA samples, CT was used as a binary classifier for CWD status. Receiver Operating Characteristic (ROC) analysis was performed to compare the CT values (obtained from DNA extracted from each control tissue homogenate) with the CWD status of those same homogenates, as determined by RT-QuIC.

Every hour during the RT-QuIC run was considered as a potential assay duration cut-off and its corresponding sensitivity was plotted against false positive rate (1-specificity). The optimal assay durations were defined as the CT with the highest Youden index (sensitivity + specificity – 1) [30]. ROC curve calculations were performed with the RStudio ROCR package [31].

#### Detection of prion seeding activity in DNA samples

RT-QuIC assays were used to determine whether DNA extracted from the archived obex (n=48) and RPLN (n=37) tissue homogenates contained any prion seeding activity. DNA samples were tested in quadruplicate. If all four replicates showed consistent seeding activity, no further testing was performed. If results were inconsistent, the sample was retested with four additional technical replicates, for a total of eight. Each DNA sample was classified as positive if at least 4 out of 8 replicates showed a CT value within the optimal assay duration; otherwise, it was classified as negative. To calculate precision, accuracy, specificity, and sensitivity of the RT-QuIC assay, the seeding activity of the DNA samples was compared to the CWD status of the source tissue homogenates, which was determined by ELISA screening and IHC confirmation. Sensitivity was defined as the ability of RT-QuIC to detect prion seeding activity in DNA extracted from a CWD+ tissue homogenate; specificity was defined as the ability of RT-QuIC to identify no prion seeding activity in DNA extracted from a CWD-tissue homogenate.

#### Evaluation of the assay duration time

To evaluate the impact of the assay duration time on the detection of prion seeding activity in DNA samples, CT values collected within the optimal or 65-h assay duration time were compared based on DNA extracted from the archived tissue homogenates. The comparison was performed at a replicate level with McNemar’s test [32] using the optimal assay duration time classification as the expected results and the 65-h assay duration time classification as the observed results.

## Results

Our results are organized according to the independent analyses conducted by each research team (MNPRO and CFIA).

### MNPRO

Average DNA yield from deer tissue samples differed significantly between extraction kits: 200 ng/µL with the DNeasy kit versus 135 ng/µL with the MagAttract kit (p < 0.0001). With the MagAttract kit, no significant difference was observed between DNA concentrations from DNA samples that tested positive versus negative for CWD seeding activity (139 vs. 129 ng/µL; p = 0.878). In contrast, with the DNeasy kit, DNA concentrations were significantly higher in samples that tested RT-QuIC positive compared to negative (267 vs. 105 ng/µL; p = 0.0001). Across both kits and species, DNA eluates testing positive for seeding activity ranged from 0.3 to 1,620 ng/µL, while those testing negative ranged from <0.1 to 590 ng/µL (Tables 1, A1–A3).

**Table 1.** DNA quantification (minimum - maximum (average)) for all DNA-extracted samples in MNPRO experiments.

Deer Samples
|  | **DNeasy DNA (ng/μL)** | **MagAttract DNA (ng/μL)** |
| --- | --- | --- |
| **All DNA tested** | 0.2 - 1620 (200) | 0.2 - 742 (135) |
| **DNA that tested positive by RT-QuIC** | 0.3 - 1620 (267) | 0.3 - 742 (139) |
| **DNA that tested negative by RT-QuIC** | <0.1 - 590 (105) | 0.2 - 512 (129) |

Hamster Samples
|  | **DNeasy DNA (ng/μL)** |
| --- | --- |
| **All DNA tested** | 4.0 - 12 (7.4) |
| **DNA that tested positive by RT-QuIC** | 4.6 - 11 (7.9) |
| **DNA that tested negative by RT-QuIC** | 4.0 - 12 (6.9) |

All 23 tissues from CWD-negative animals, along with their paired DNA eluates from both extraction kits, tested negative by RT-QuIC. Among the 53 tissues from CWD-positive animals, 32 were RT-QuIC positive in both the tissue and the paired DNA extracts from both kits. Twelve samples from CWD-positive animals were negative across tissue and DNA eluates from both kits; this group included all eight muscle samples. In addition, two samples were RT-QuIC positive in tissue but negative in both DNA eluates, while three were positive in tissue and in only one of the two DNA extracts. Notably, four samples (brain stem, ear, and two salivary glands) were RT-QuIC positive only in the DNA eluates from both extraction kits, but not in the corresponding tissue homogenates (Table A2).

Agreement between each animal’s CWD status (determined by ELISA/IHC of RPLN) and the RT-QuIC seeding activity of DNA eluates was evaluated using Cohen’s kappa (κ). Moderate agreement [33] was observed for both DNA extraction methods: DNeasy eluates (κ = 0.583; SE = 0.084; 95% CI: 0.418 - 0.748) and MagAttract eluates (κ = 0.605; SE = 0.084; 95% CI: 0.441 - 0.770). Because perfect agreement is not expected between every individual tissue and the overall CWD status of the animal, particularly in early-stage infections, we also compared the seeding activity of DNA eluates with that of their corresponding tissue homogenates. This analysis showed substantial agreement for DNeasy eluates (κ = 0.789; SE = 0.070; 95% CI: 0.651 - 0.927) and almost perfect agreement for MagAttract eluates (κ = 0.816; SE = 0.066; 95% CI: 0.686 - 0.946; Table 2). There was 100% agreement between the experimental status of hamsters and seeding activity in their DNA eluates and the corresponding tissue homogenates.

**Table 2.** Cohen’s kappa data for MNPRO WTD tissue and DNA eluates. Panel A compares the seeding activity of DNA eluates to the CWD status of the animals. Panel B compares the seeding activity of the DNA eluates to that of the source tissue. DNE=DNeasy. MA=MagAttract.

| **A.** | **DNE DNA that tested positive by RT-QuIC** | **DNE DNA that tested negative by RT-QuIC** | **Total** | **Kappa** | **Kappa interpretation** | **SE of kappa** | **95% CI of kappa** |
| --- | --- | --- | --- | --- | --- | --- | --- |
| Tissue obtained from a CWD-positive WTD | 37 | 16 | 53 | 0.583 | moderate agreement | 0.084 | 0.418 - 0.748 |
| Tissue obtained from a CWD-negative WTD | 0 | 23 | 23 |  |  |  |  |
| **Total** | 37 | 39 | 76 |  |  |  |  |
|  | **MA DNA that tested positive by RT-QuIC** | **MA DNA that tested negative by RT-QuIC** | **Total** | **Kappa** | **Kappa interpretation** | **SE of kappa** | **95% CI of kappa** |
| Tissue obtained from a CWD-positive WTD | 38 | 15 | 53 | 0.605 | moderate agreement | 0.084 | 0.441 - 0.770 |
| Tissue obtained from a CWD-negative WTD | 0 | 23 | 23 |  |  |  |  |
| **Total** | 38 | 38 | 76 |  |  |  |  |

| **B.** | **DNE DNA that tested positive by RT-QuIC** | **DNE DNA that tested negative by RT-QuIC** | **Total** | **Kappa** | **Kappa interpretation** | **SE of kappa** | **95% CI of kappa** |
| --- | --- | --- | --- | --- | --- | --- | --- |
| Tissue homogenate that tested positive by RT-QuIC | 33 | 4 | 37 | 0.789 | substantial agreement | 0.07 | 0.651 - 0.927 |
| Tissue homogenate that tested negative by RT-QuIC | 4 | 35 | 39 |  |  |  |  |
| **Total** | 37 | 39 | 76 |  |  |  |  |
|  | **MA DNA that tested positive by RT-QuIC** | **MA DNA that tested negative by RT-QuIC** | **Total** | **Kappa** | **Kappa interpretation** | **SE of kappa** | **95% CI of kappa** |
| Tissue homogenate that tested positive by RT-QuIC | 34 | 3 | 37 | 0.816 | almost perfect agreement | 0.066 | 0.686 - 0.946 |
| Tissue homogenate that tested negative by RT-QuIC | 4 | 35 | 39 |  |  |  |  |
| **Total** | 38 | 38 | 76 |  |  |  |  |

Endpoint titration studies on nine positive samples revealed a further endpoint with higher DNA concentrations (Figure 1 and Table A4). In contrast, the endpoint for paired tissue samples remained relatively stable. A negative sample (not shown in Figure 1) was also tested in a serial dilution and exhibited no seeding activity in either the tissue or the extracted DNA (Table A4).

**Figure 1.**
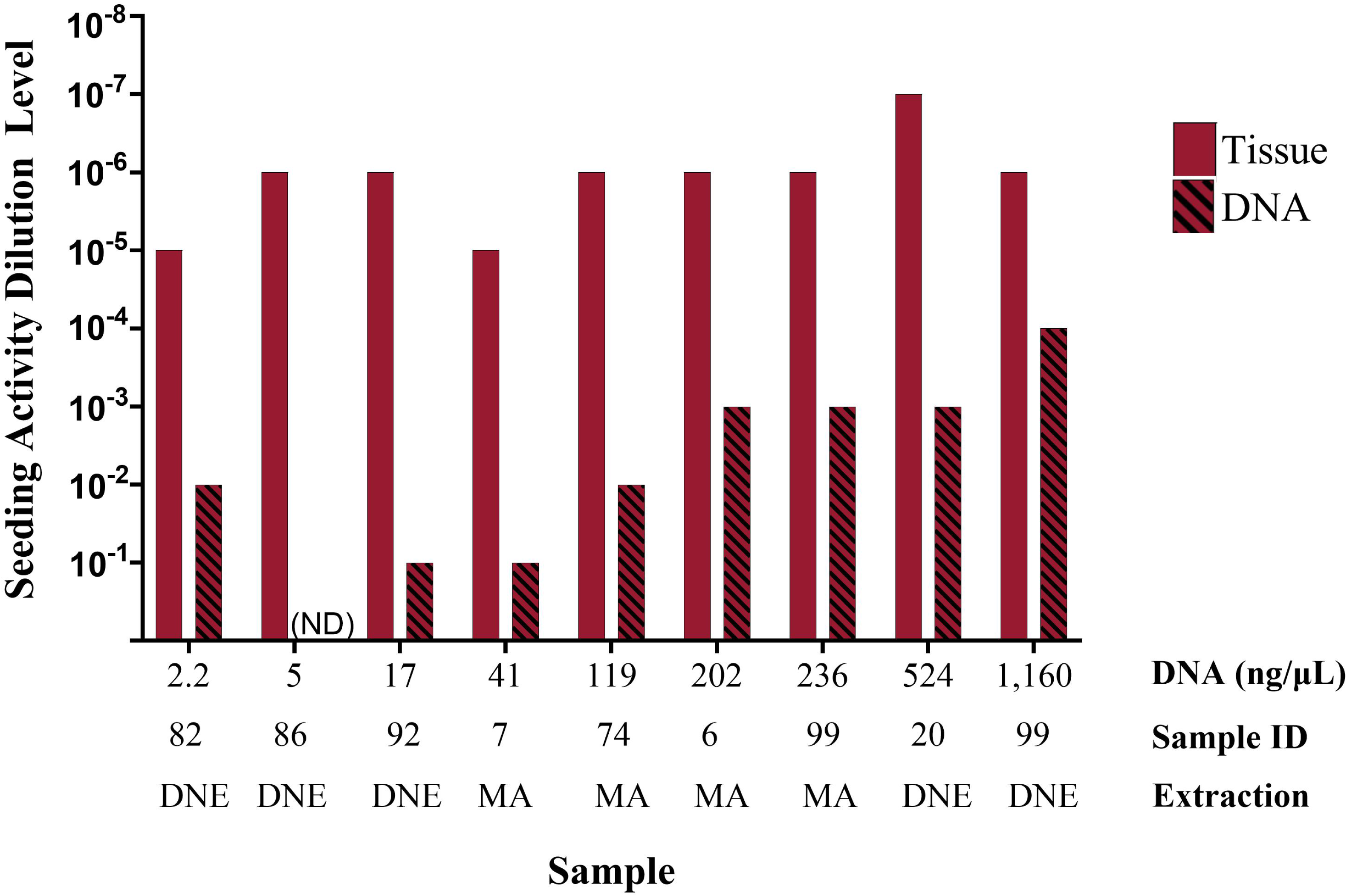
RT-QuIC endpoint titration experiment of CWD-positive tissue and matched DNA for a subset of positive samples with increasing DNA concentrations. Bar height indicates the final dilution where statistically significant seeding activity was observed. Additional dilution levels tested are not represented in this figure but are shown in Table A4. ND = not detected. Sample IDs correspond to the Sample IDs in Table A2. Extraction: DNE = DNeasy; MA = MagAttract.

Simple linear regression was performed to determine whether sample DNA concentration predicted the endpoint for extracted DNA. The analysis revealed a statistically significant relationship (R^2^ = 0.66, F (1, 6) = 11.77, p-value = 0.014), indicating that DNA concentration influences the endpoint titration of extracted DNA when assessing seeding activity by RT-QuIC.

Similarly, simple linear regression was used to examine whether DNA concentration predicted the endpoint titration for tissue samples. However, this regression was not statistically significant (R^2^ = 0.11, F (1, 7) = 0.84, p-value = 0.399), suggesting that DNA concentration does not predict the limit-of-detection of tissue samples when testing for seeding activity by RT-QuIC.

### CFIA

#### CWD status of the control tissue homogenates

To optimize the RT-QuIC assay for detecting prion seeding activity in DNA samples, control tissue homogenates were prepared from serial dilutions of CWD+ in CWD-obex and RPLN tissue homogenates. Figure 2 shows the CT values of the control tissue homogenates using our previously established optimal assay duration time of 30 and 33 h for obex and RPLN tissues, respectively [28]. The control homogenates containing CWD+ or 10^-1^ and 10^-2^ of CWD+ obex and RPLN tissue tested positive for CWD by RT-QuIC (Table 3). The control homogenates containing only CWD- or 10^-5^ to 10^-3^ CWD+ obex and RPLN tissue homogenate tested negative for CWD by RT-QuIC (Table 3).

**Figure 2.**
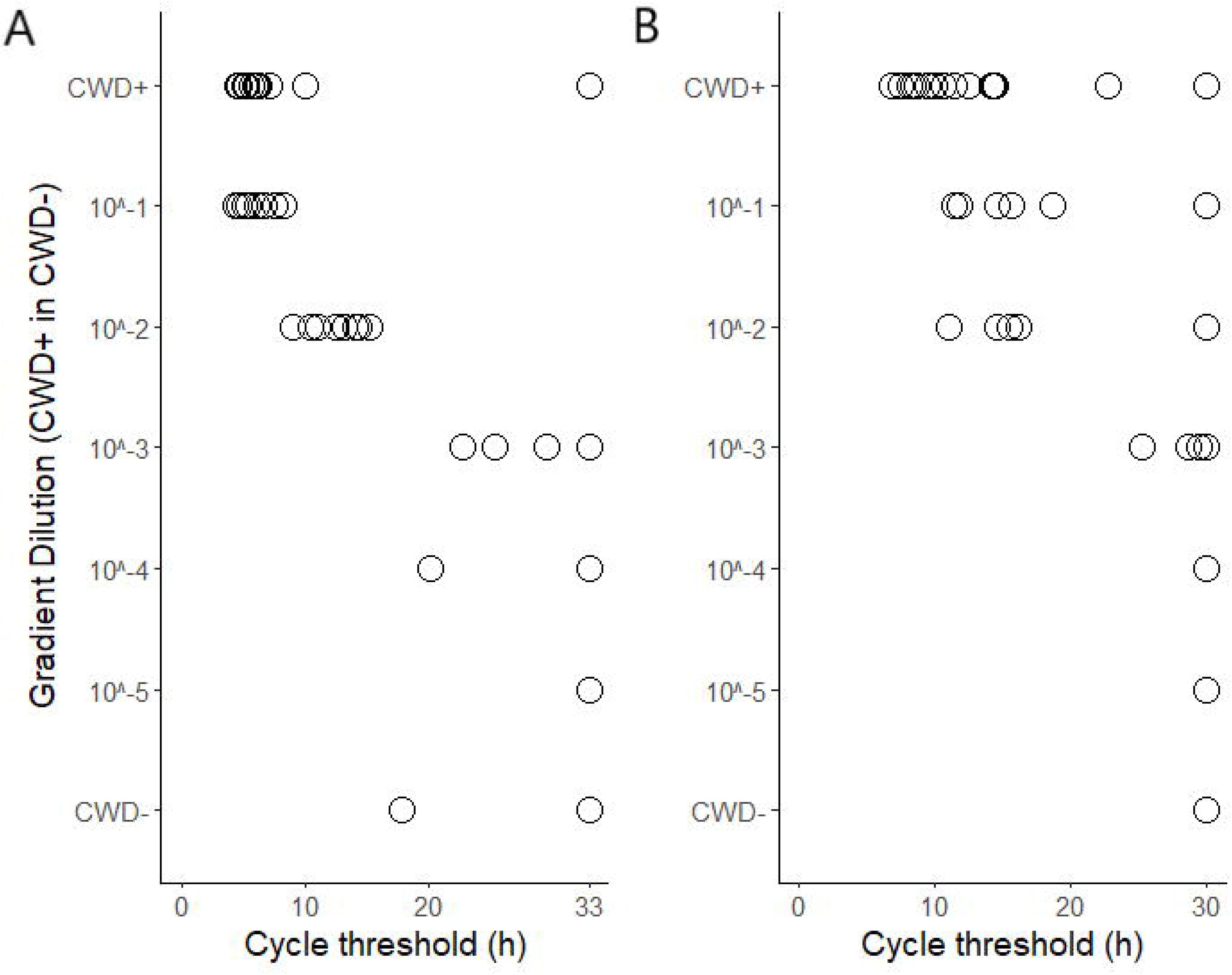
Cycle threshold values of CFIA individual RT-QuIC reactions on CWD+, CWD-, and 10-fold serial dilutions of CWD+ in CWD-obex and retropharyngeal lymph node (RPLN) tissue homogenates within the 33-h and 30-h assay time for obex (A) and RPLN (B), respectively.

**Table 3.**
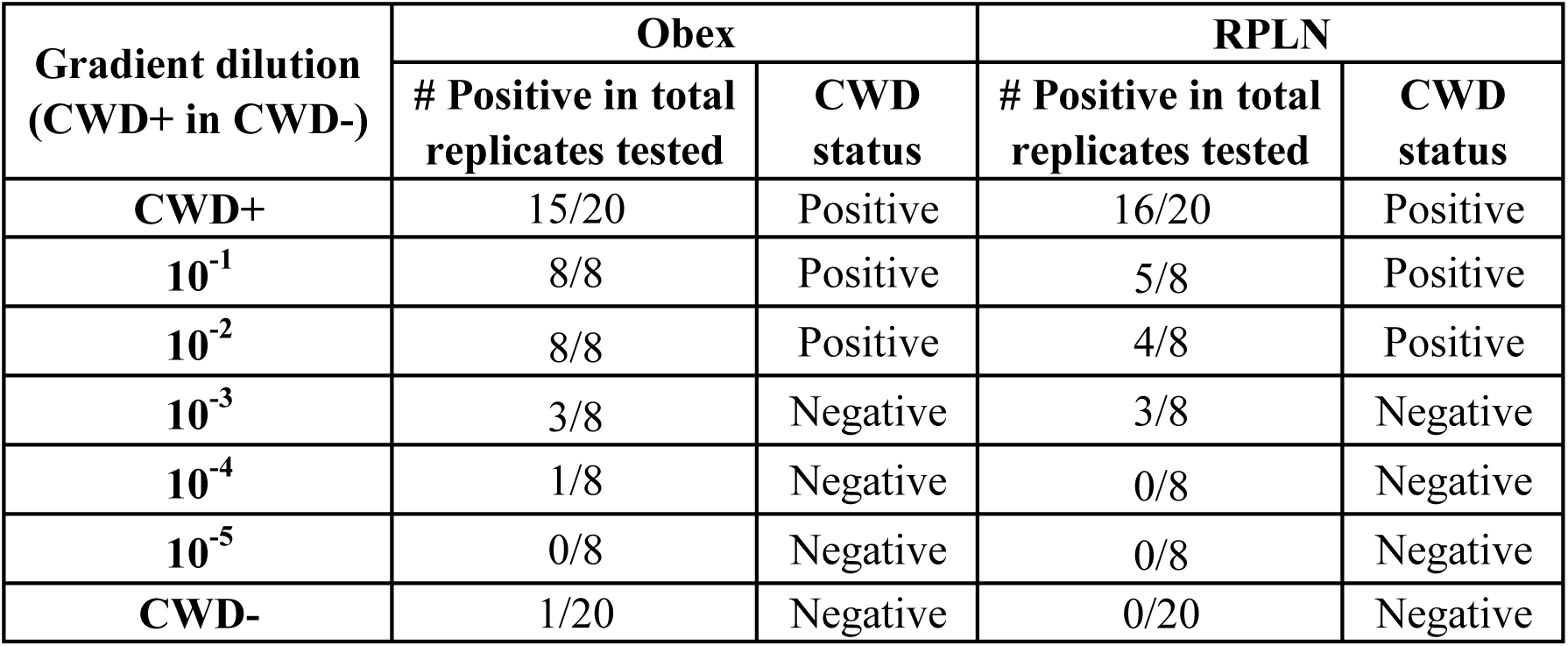
Chronic wasting disease (CWD) status^1^ of the control (obex and RPLN) tissue homogenates^2^ by RT-QuIC. ^1^A tissue homogenate was positive by RT-QuIC when ThT signal of at least half of its replicates crossed the threshold within the 33-h and 30-h assay time for obex and RPLN, respectively; otherwise, it was negative. ^2^Tissue homogenates were mixtures of CWD+ and CWD-tissue homogenate in various ratios, each containing 0.01% (w/v) of tissue specimens.

| Gradient dilution<br>(CWD+ in CWD-) | Obex |  | RPLN |  |
| --- | --- | --- | --- | --- |
|  | # Positive in total<br>replicates tested | CWD<br>status | # Positive in total<br>replicates tested | CWD<br>status |
| **CWD+** | 15/20 | Positive | 16/20 | Positive |
| **10<sup>-1</sup>** | 8/8 | Positive | 5/8 | Positive |
| **10<sup>-2</sup>** | 8/8 | Positive | 4/8 | Positive |
| **10<sup>-3</sup>** | 3/8 | Negative | 3/8 | Negative |
| **10<sup>-4</sup>** | 1/8 | Negative | 0/8 | Negative |
| **10<sup>-5</sup>** | 0/8 | Negative | 0/8 | Negative |
| **CWD-** | 1/20 | Negative | 0/20 | Negative |

#### Optimal RT-QuIC assay duration for detection of prion seeding activity in DNA samples

DNA extracted from the above control tissue homogenates had a mean concentration of 31.00 ng/µL (Table B1). To minimize the loss of potential prion protein, these DNA samples underwent RT-QuIC assays without normalization according to their concentrations. Their CT values from RT-QuIC are shown in Figure 3A. There was negligible correlation (Spearman correlation efficiency, r = −0.265) between the concentration and the corresponding CT value of the DNA samples (Fig. B1). A global ROC curve was constructed using the CT values of the DNA samples against the CWD status of the corresponding control tissue homogenates determined by RT-QuIC (Figure 3.B). Based on the ROC analysis, 30 hours was identified as the optimal assay duration, producing the fewest false positives and the highest sensitivity. The observed area under the ROC curve (AUC) was 0.731, indicating that the optimal assay duration time provides a decent classifying power for detecting prion seeding activity in DNA samples.

**Figure 3.**
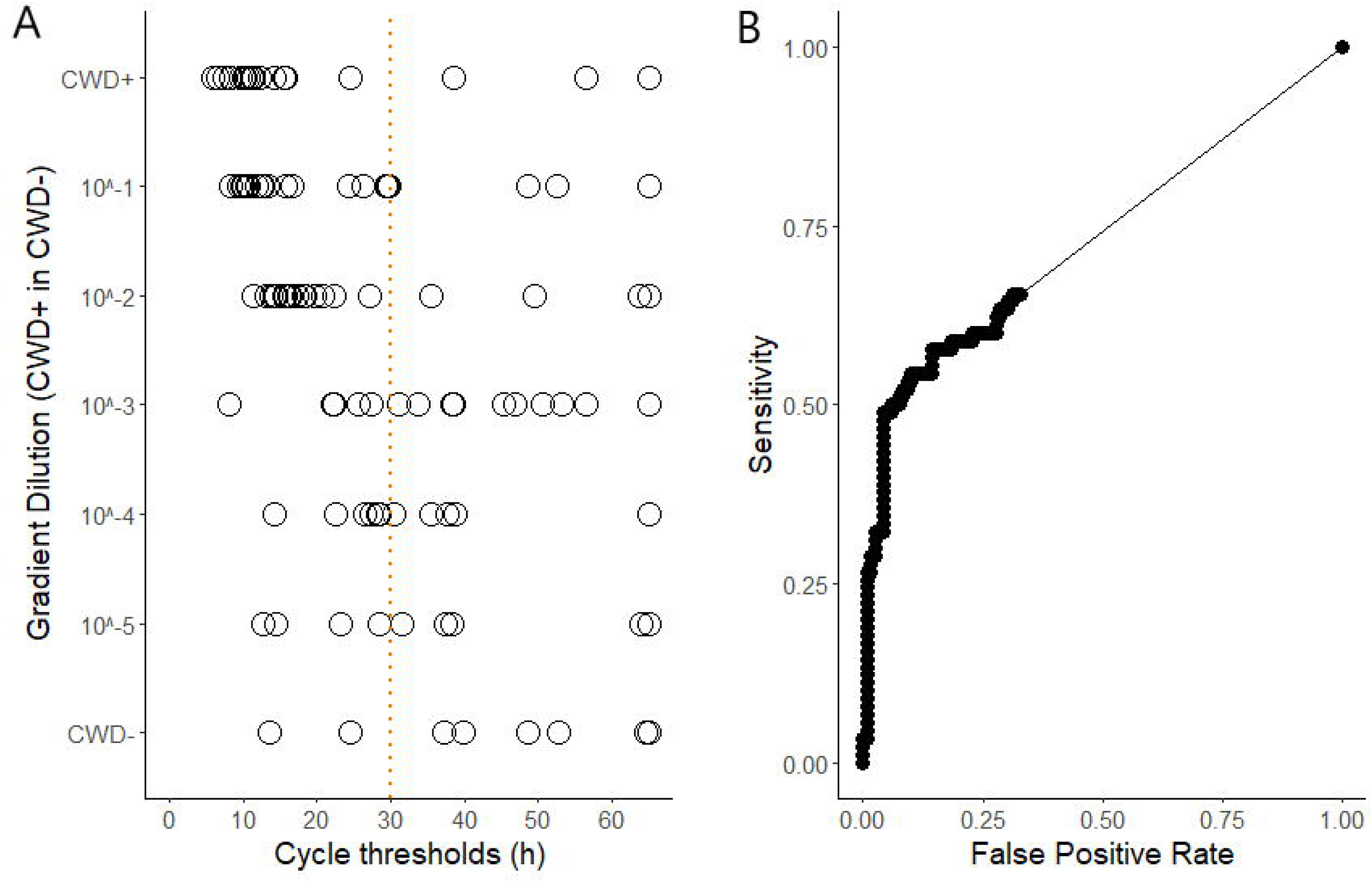
Cycle threshold (CT) values of individual RT-QuIC reactions on DNA extracted from CWD+, CWD-, and 10-fold serial dilutions of CWD+ in CWD-obex and RPLN tissue homogenates using a 65-h assay time (A). A global receiver operating characteristic (ROC) curve (B) was constructed using the CT values on DNA against the CWD status of the corresponding tissue homogenates determined by RT-QuIC. The dashed line in panel A indicates the optimal assay time identified using the ROC curve in panel B.

#### Prion seeding activity in DNA samples from archived tissue specimens

DNA extracted from 88 archived tissue specimens was tested using RT-QuIC to assess the presence of prion seeding activity. The samples included 48 obex and 40 RPLN specimens. Of these, 28 obex and 25 RPLN samples had been previously diagnosed as CWD-positive by ELISA and confirmed by IHC. RT-QuIC detected prion seeding activity in 19 obex and 20 RPLN DNA samples from the CWD-positive group, while 9 obex and 5 RPLN DNA samples tested negative (Table 4). Among the CWD-negative specimens, 1 obex and 1 RPLN DNA sample tested positive, while 19 obex and 14 RPLN DNA samples tested negative (Table 4).

**Table 4.** Prion seeding activity in DNA extracted from archived obex and retropharyngeal lymph node (RPLN) tissue specimens by RT-QuIC. ^1^DNA was considered positive for prion seeding activity by RT-QuIC when the ThT signal of at least 4/4 or 4/8 replicates crossed the threshold within the 30-h assay time.

|  |  |  | **Prion seeding activity in**<br>**DNA by RT-QuIC<sup>1</sup>** |  |
| --- | --- | --- | --- | --- |
|  |  |  | **Positive** | **Negative** |
| **CWD status of**<br>**archived tissue**<br>**homogenates by**<br>**ELISA** | Obex | Positive | 19 | 9 |
|  |  | Negative | 1 | 19 |
|  | RLN | Positive | 20 | 5 |
|  |  | Negative | 1 | 14 |

#### Evaluation of the optimal RT-QuIC assay duration time

Figure 4 and Table B3 show RT-QuIC CT values from DNA samples of 88 archived tissue homogenates. With both 30-h and 65-h assay times, most positive samples had CT values below 30 hours. More high CT values were observed in negative samples after 65 h than after 30 h. The 30-h assay showed lower sensitivity (74% vs. 87%) but higher specificity (94% vs. 74%) compared with the 65-h assay. This performance difference was significant by McNemar’s test (Table 5).

**Figure 4.**
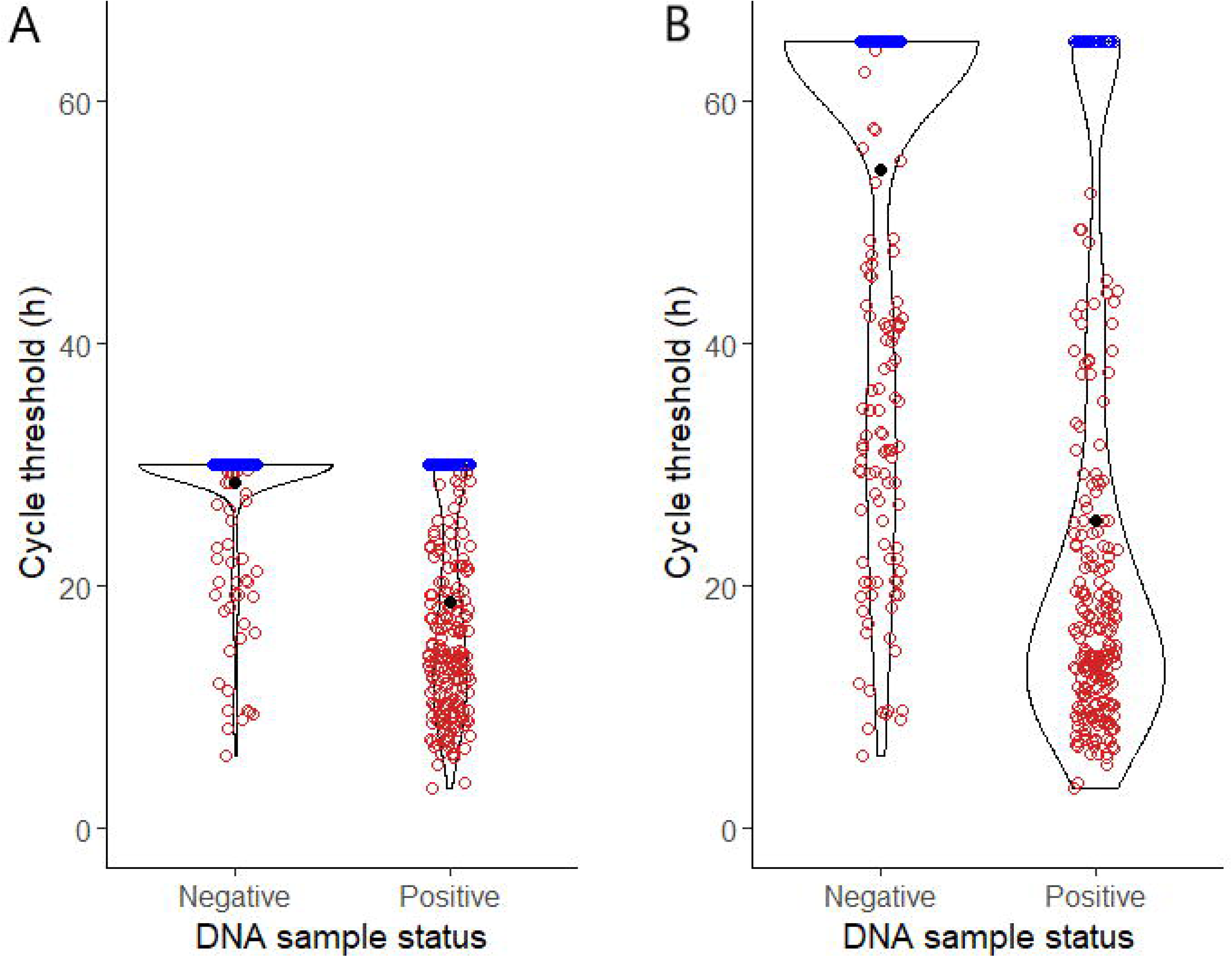
Cycle threshold values of individual RT-QuIC reactions on DNA extracted from archived WTD tissue homogenates. DNA samples were determined positive or negative for prion seeding activity by RT-QuIC using the 30-h (A) and 65-h (B) assay time. ThT signals of the reactions in red crossed the threshold within the corresponding assay time, whereas those of the reactions in blue did not.

**Table 5.** Comparison of RT-QuIC for the detection of prion seeding activity in DNA samples using 30-h and 65-h assay times. ^1^True positive (TP) represents the number of DNA extracted from CWD+ tissue homogenates tested positive for prion seeding activity by RT-QuIC. ^2^True negative (TN) represents the number of DNA extracted from CWD-tissue homogenates tested negative for prion seeding activity by RT-QuIC ^3^False positive (FP) represents the number of DNA extracted from CWD-tissue homogenates tested positive for prion seeding activity by RT-QuIC. ^4^False negative (FN) represents the number of DNA extracted from CWD+ tissue homogenates tested negative for prion seeding activity by RT-QuIC. ^5^Sensitivity = TP / (TP + FN) ^6^Specificity = TN / (TN + FP).

| **Assay time** | **n** | **TP<sup>1</sup>** | **TN<sup>2</sup>** | **FP<sup>3</sup>** | **FN<sup>4</sup>** | **Sensitivity<sup>5</sup>** | **Specificity<sup>6</sup>** | **McNemar's Statistic** | ***p* -value** |
| --- | --- | --- | --- | --- | --- | --- | --- | --- | --- |
| **30-h** | 88 | 39 | 33 | 2 | 14 | 74% | 94% | 12.071 | <0.001 |
| **65-h** | 88 | 46 | 26 | 9 | 7 | 87% | 74% |  |  |

## Discussion

Our combined findings provide strong and convergent evidence that DNA extracts from PrP^Sc^-positive tissues can retain substantial RT-QuIC prion seeding activity. This observation introduces a previously unrecognized biosafety and biosecurity concern for laboratories handling nucleic acids from prion-positive sources.

In this work, both laboratories independently observed robust detection of PrP^Sc^ in DNA eluates. MNPRO identified seeding activity across a wide range of DNA concentrations, from 0.3 ng/μl to over 1,600 ng/μl (Table 1), demonstrating substantial to almost perfect agreement between the RT-QuIC seeding activity of DNA eluates and their source tissue homogenates (Table 2). DNA derived from WTD tissues, including ear, lymph node, salivary gland, third eyelid, tongue, and tonsil, tested positive by RT-QuIC, corroborating previous findings from IHC, ELISA, PMCA, and RT-QuIC performed on homogenates from these tissue types [33–37].

A ROC analysis was used by the CFIA laboratory to refine assay conditions, leading to a reduced RT-QuIC runtime of 30 hours. This optimized protocol yielded positive results in 39 of 53 DNA samples from CWD-positive tissues and only 2 positives among 35 CWD-negative controls, resulting in 74% sensitivity and 94% specificity. While this abbreviated assay displayed slightly reduced sensitivity compared to a 65-hour protocol, it offered improvements in both specificity and sample throughput. Together, these results demonstrate that PrP^Sc^ can co-purify with DNA during standard extraction procedures, regardless of the extraction chemistry used (filter-based or magnetic bead-based) and across a variety of tissue types.

Several mechanistic factors likely contribute to the co-extraction of PrP^Sc^ in DNA extraction workflows. Many commercial DNA extraction kits incorporate PK digestion steps that would degrade PrP^C^, likely enriching for protease-resistant PrP^Sc^, resulting in its presence in the final eluate. In addition, PrP^C^ has been shown to bind directly to DNA [39], and in vitro studies have demonstrated that polyanionic nucleic acids can facilitate the conformational conversion of PrP^C^ to PrP^Sc^ [11]. It is plausible that DNA-protein complexes formed during tissue lysis protect PrP^Sc^ from subsequent wash or filter steps, allowing it to remain in the purified DNA fraction. Our data further indicate that residual PrP^Sc^, rather than the DNA itself, accounts for the RT-QuIC seeding activity, as no seeding activity was detected in MNPRO DNA eluates from 23 CWD-negative tissues, despite DNA concentrations averaging 128 ng/µL (range 0.2–590 ng/µL; Table A1).

While CFIA observed no correlation between DNA concentration and RT-QuIC CT values in DNA eluates of spiked control samples, MNPRO found that DNA concentration influenced the seeding activity observed during endpoint titration of DNA extracts from CWD-positive tissues. Both CFIA and MNPRO reported that when DNA concentrations are below 60 ng/µL, seeding activity becomes inconsistent or random. However, at concentrations above this threshold, higher DNA levels appear to enhance seeding activity.

The presence of PrP^Sc^ in DNA extracts introduces a layer of biosafety risk that must be considered when handling nucleic acids derived from PrP^Sc^-positive materials. Laboratory personnel working with DNA samples presumed to be non-infectious may unknowingly be exposed to potentially infectious material. Routine molecular workflows involving these DNA extracts could also lead to cross-contamination of laboratory surfaces, equipment, and reagents. Due to these risks, handling DNA from known or suspected TSE-positive sources under containment conditions should be considered. This includes using prion-effective decontamination procedures and treating DNA preparations as potentially infectious unless proven otherwise.

Several key questions remain unanswered. We have not evaluated the in vivo infectivity of DNA eluates. Planned mouse bioassays will be essential for determining infectivity, quantifying infectious titers by relevant routes of infection, and refining the biosafety classification of DNA extracted from prion-infected tissues. Additionally, systematic comparisons of different lysis buffers, binding media, and wash conditions could help identify protocols that minimize PrP^Sc^ carryover without compromising nucleic acid integrity. Standardization of RT-QuIC protocols, including harmonized cutoffs, reaction times, and reference materials, will also be necessary to enable reproducible inter-laboratory comparisons and broader diagnostic applications.

Both participating laboratories independently reached the same fundamental conclusion: standard DNA extraction procedures do not guarantee the removal of PrP^Sc^. Until the infectivity of DNA extracts from TSE-positive tissues is fully characterized, our findings suggest that all nucleic-acid preparations derived from such materials should be handled as potentially infectious. Enhanced biosafety measures, validated workflows, and rigorous decontamination protocols will be essential to protect laboratory personnel, prevent cross-contamination, and ensure the continued safe advancement of genetic and molecular research in prion diseases.

## Acknowledgments

This project was partially funded by the Minnesota Environment and Natural Resources Trust Fund, as recommended by the Legislative-Citizen Commission on Minnesota Resources (LCCMR). We would like to thank the Creighton University Animal Resource Facility for outstanding animal care. We thank the entire MNPRO team, especially Suzanne Stone, for her lab management. CFIA’s study was supported by the Canadian Food Inspection Agency Research Fund N000482. We thank the current and previous staff of the CFIA – Operations Unit and the CFIA – Transmissible Spongiform Encephalopathy Unit for their contribution in the supply, processing, and testing of the deer samples.

We thank the Minnesota Department of Natural Resources for providing access to biological samples from free-ranging WTD that were collected through state-funded CWD surveillance programs.

## References

[1] S. B. Prusiner, D. F. Groth, C. Bildstein, F. R. Masiarz, M. P. McKinley, and S. P. Cochran, “Electrophoretic properties of the scrapie agent in agarose gels,” Proc. Natl. Acad. Sci. U. S. A., vol. 77, no. 5, pp. 2984–2988, May 1980, doi: 10.1073/pnas.77.5.2984.

[2] M. P. McKinley, D. C. Bolton, and S. B. Prusiner, “A protease-resistant protein is a structural component of the scrapie prion,” Cell, vol. 35, no. 1, pp. 57–62, Nov. 1983, doi: 10.1016/0092-8674(83)90207-6.

[3] P. Brown, A. Wolff, and D. C. Gajdusek, “A simple and effective method for inactivating virus infectivity in formalin-fixed tissue samples from patients with Creutzfeldt-Jakob disease,” Neurology, vol. 40, no. 6, pp. 887–887, June 1990, doi: 10.1212/WNL.40.6.887.

[4] P. Brown, R. G. Rohwer, E. M. Green, and D. C. Gajdusek, “Effect of chemicals, heat, and histopathologic processing on high-infectivity hamster-adapted scrapie virus,” J. Infect. Dis., vol. 145, no. 5, pp. 683–687, May 1982, doi: 10.1093/infdis/145.2.683.

[5] J.-P. Brandel et al., “Variant Creutzfeldt-Jakob Disease Diagnosed 7.5 Years after Occupational Exposure,” N. Engl. J. Med., vol. 383, no. 1, pp. 83–85, July 2020, doi: 10.1056/NEJMc2000687.

[6] B. Casassus, “France halts prion research amid safety concerns,” Science, vol. 373, no. 6554, pp. 475–476, July 2021, doi: 10.1126/science.373.6554.475.

[7] C. J. Silva, M. L. Erickson Beltran, and J. R. Requena, “Comparing the Extent of Methionine Oxidation in the Prion and Native Conformations of PrP,” ACS Omega, vol. 10, no. 1, pp. 1320–1330, Jan. 2025, doi: 10.1021/acsomega.4c08892.

[8] A. M. Thackray, L. Hopkins, and R. Bujdoso, “Proteinase K-sensitive disease-associated ovine prion protein revealed by conformation-dependent immunoassay,” Biochem. J., vol. 401, no. Pt 2, pp. 475–483, Jan. 2007, doi: 10.1042/BJ20061264.

[9] M. A. Pastrana et al., “Isolation and characterization of a proteinase K-sensitive PrPSc fraction,” Biochemistry, vol. 45, no. 51, pp. 15710–15717, Dec. 2006, doi: 10.1021/bi0615442.

[10] A. Brun et al., “Proteinase K enhanced immunoreactivity of the prion protein-specific monoclonal antibody 2A11,” Neurosci. Res., vol. 48, no. 1, pp. 75–83, Jan. 2004, doi: 10.1016/j.neures.2003.09.004.

[11] Y. Cordeiro et al., “DNA Converts Cellular Prion Protein into the β-Sheet Conformation and Inhibits Prion Peptide Aggregation*,” J. Biol. Chem., vol. 276, no. 52, pp. 49400–49409, Dec. 2001, doi: 10.1074/jbc.M106707200.

[12] C. Gabus et al., “The prion protein has RNA binding and chaperoning properties characteristic of nucleocapsid protein NCP7 of HIV-1,” J. Biol. Chem., vol. 276, no. 22, pp. 19301–19309, June 2001, doi: 10.1074/jbc.M009754200.

[13] N. R. Deleault, R. W. Lucassen, and S. Supattapone, “RNA molecules stimulate prion protein conversion,” Nature, vol. 425, no. 6959, pp. 717–720, Oct. 2003, doi: 10.1038/nature01979.

[14] J. C. Geoghegan et al., “Selective incorporation of polyanionic molecules into hamster prions,” J. Biol. Chem., vol. 282, no. 50, pp. 36341–36353, Dec. 2007, doi: 10.1074/jbc.M704447200.

[15] W.-Q. Zou, J. Zheng, D. M. Gray, P. Gambetti, and S. G. Chen, “Antibody to DNA detects scrapie but not normal prion protein,” Proc. Natl. Acad. Sci., vol. 101, no. 5, pp. 1380–1385, Feb. 2004, doi: 10.1073/pnas.0307825100.

[16] P. K. Nandi and J.-C. Nicole, “Nucleic acid and prion protein interaction produces spherical amyloids which can function in vivo as coats of spongiform encephalopathy agent,” J. Mol. Biol., vol. 344, no. 3, pp. 827–837, Nov. 2004, doi: 10.1016/j.jmb.2004.09.080.

[17] J. A. Blow, D. J. Dohm, D. L. Negley, and C. N. Mores, “Virus inactivation by nucleic acid extraction reagents,” J. Virol. Methods, vol. 119, no. 2, pp. 195–198, Aug. 2004, doi: 10.1016/j.jviromet.2004.03.015.

[18] J. Collinge, M. S. Palmer, and A. J. Dryden, “Genetic predisposition to iatrogenic Creutzfeldt-Jakob disease,” The Lancet, vol. 337, no. 8755, pp. 1441–1442, June 1991, doi: 10.1016/0140-6736(91)93128-V.

[19] S. Mead et al., “Genetic risk factors for variant Creutzfeldt–Jakob disease: a genome-wide association study,” Lancet Neurol., vol. 8, no. 1, pp. 57–66, Jan. 2009, doi: 10.1016/S1474-4422(08)70265-5.

[20] H. Geldermann, H. He, P. Bobal, H. Bartenschlager, and S. Preuß, “Comparison of DNA variants in the PRNP and NF1 regions between bovine spongiform encephalopathy and control cattle,” Anim. Genet., vol. 37, no. 5, pp. 469–474, 2006, doi: 10.1111/j.1365-2052.2006.01519.x.

[21] J. A. Richt et al., “Identification and Characterization of two Bovine Spongiform Encephalopathy cases Diagnosed in the United States,” J. Vet. Diagn. Invest., vol. 19, no. 2, pp. 142–154, Mar. 2007, doi: 10.1177/104063870701900202.

[22] G. C. Saunders, S. Cawthraw, S. J. Mountjoy, J. Hope, and O. Windl, “PrP genotypes of atypical scrapie cases in Great Britain,” J. Gen. Virol., vol. 87, no. 11, pp. 3141–3149, 2006, doi: 10.1099/vir.0.81779-0.

[23] N. J. Haley et al., “Management of chronic wasting disease in ranched elk: conclusions from a longitudinal three-year study,” Prion, vol. 14, no. 1, pp. 76–87, Jan. 2020, doi: 10.1080/19336896.2020.1724754.

[24] C. I. Cullingham, E. H. Merrill, M. J. Pybus, T. K. Bollinger, G. A. Wilson, and D. W. Coltman, “Broad and fine-scale genetic analysis of white-tailed deer populations: estimating the relative risk of chronic wasting disease spread,” Evol. Appl., vol. 4, no. 1, pp. 116–131, Jan. 2011, doi: 10.1111/j.1752-4571.2010.00142.x.

[25] A. L. Brandt, M. L. Green, Y. Ishida, A. L. Roca, J. Novakofski, and N. E. Mateus-Pinilla, “Influence of the geographic distribution of prion protein gene sequence variation on patterns of chronic wasting disease spread in white-tailed deer (Odocoileus virginianus),” Prion, vol. 12, no. 3–4, pp. 204–215, July 2018, doi: 10.1080/19336896.2018.1474671.

[26] A. J. Block et al., “Efficient interspecies transmission of synthetic prions,” PLoS Pathog., vol. 17, no. 7, p. e1009765, July 2021, doi: 10.1371/journal.ppat.1009765.

[27] M. D. Schwabenlander et al., “Comparison of chronic wasting disease detection methods and procedures: Implications for free-ranging white-tailed deer (Odocoileus virginianus) surveillance and management,” J. Wildl. Dis., vol. 58, no. 1, pp. 50–62, Jan. 2022, doi: 10.7589/JWD-D-21-00033.

[28] G. Yilmaz et al., “Optimization of RT-QuIC Assay Duration for Screening Chronic Wasting Disease in White-Tailed Deer,” Vet. Sci., vol. 11, no. 2, p. 60, Feb. 2024, doi: 10.3390/vetsci11020060.

[29] H.-J. Sohn et al., “Experimental oral transmission of chronic wasting disease to sika deer (Cervus nippon),” Prion, vol. 14, no. 1, pp. 271–277, Dec. 2020, doi: 10.1080/19336896.2020.1857038.

[30] I. Unal, “Defining an Optimal Cut-Point Value in ROC Analysis: An Alternative Approach,” Comput. Math. Methods Med., vol. 2017, p. 3762651, 2017, doi: 10.1155/2017/3762651.

[31] T. Sing, O. Sander, N. Beerenwinkel, and T. Lengauer, “ROCR: visualizing classifier performance in R,” Bioinforma. Oxf. Engl., vol. 21, no. 20, pp. 3940–3941, Oct. 2005, doi: 10.1093/bioinformatics/bti623.

[32] A. Agresti, Categorical Data Analysis, 3rd ed. Wiley, 2013. Accessed: July 23, 2025. [Online]. Available: https://www.wiley.com/en-gb/Categorical+Data+Analysis%%2C+3rd+Edition-p-9780470463635

[33] J. R. Landis and G. G. Koch, “The Measurement of Observer Agreement for Categorical Data,” Biometrics, vol. 33, no. 1, pp. 159–174, 1977, doi: 10.2307/2529310.

[34] N. J. Haley, C. K. Mathiason, S. Carver, M. Zabel, G. C. Telling, and E. A. Hoover, “Detection of Chronic Wasting Disease Prions in Salivary, Urinary, and Intestinal Tissues of Deer: Potential Mechanisms of Prion Shedding and Transmission▿,” J. Virol., vol. 85, no. 13, pp. 6309–6318, July 2011, doi: 10.1128/JVI.00425-11.

[35] S. K. Cooper, C. E. Hoover, D. M. Henderson, N. J. Haley, C. K. Mathiason, and E. A. Hoover, “Detection of CWD in cervids by RT-QuIC assay of third eyelids,” PLoS ONE, vol. 14, no. 8, p. e0221654, Aug. 2019, doi: 10.1371/journal.pone.0221654.

[36] N. C. Ferreira et al., “Detection of chronic wasting disease in mule and white-tailed deer by RT-QuIC analysis of outer ear,” Sci. Rep., vol. 11, no. 1, p. 7702, Apr. 2021, doi: 10.1038/s41598-021-87295-8.

[37] D. A. Schneider et al., “Tonsil biopsy to detect chronic wasting disease in white-tailed deer (Odocoileus virginianus) by immunohistochemistry,” PloS One, vol. 18, no. 3, p. e0282356, 2023, doi: 10.1371/journal.pone.0282356.

[38] C. L. Holz et al., “Evaluation of Real-Time Quaking-Induced Conversion, ELISA, and Immunohistochemistry for Chronic Wasting Disease Diagnosis,” Front. Vet. Sci., vol. 8, 2022, Accessed: July 11, 2023. [Online]. Available: https://www.frontiersin.org/articles/10.3389/fvets.2021.824815

[39] A. Tandon, V. K. Subramani, K. K. Kim, and S. H. Park, “Interaction of Prion Peptides with DNA Structures,” ACS Omega, vol. 7, no. 1, pp. 176–186, Dec. 2021, doi: 10.1021/acsomega.1c04328.

