## Appendix A, Table A1 for "Prion Seeding Activity in DNA Extractions: Implications for Laboratory Biosafety"

| | | | | MagAttract DNA | | Dneasy DNA | | Tissue |
|---|---|---|---|---|---|---|---|---|
| **Animal ID** | **Animal's CWD Status (by ELISA or IHC)** | **Sample ID** | **Tissue** | **RT-QuIC Results** | **[DNA] (ng/uL)** | **RT-QuIC Results** | **[DNA] (ng/uL)** | **RT-QuIC Results** |
| 4 | - | 25 | brain | - | 232 | - | 323 | - |
| | | 26 | parotid LN | - | 504 | - | 300 | - |
| 5 | - | 27 | brain | - | 87 | - | 18 | - |
| | | 28 | parotid LN | - | 164 | - | 7.7 | - |
| 6 | - | 29 | brain | - | 302 | - | 140 | - |
| | | 30 | parotid LN | - | 61 | - | 0.7 | - |
| 7 | - | 32 | parotid LN | - | 0.2 | - | 306 | - |
| 8 | - | 33 | brain | - | 1.1 | - | 19 | - |
| | | 34 | parotid LN | - | 2.7 | - | 21 | - |
| 9 | - | 35 | brain | - | 76 | - | 28 | - |
| | | 36 | parotid LN | - | 512 | - | 590 | - |
| 10 | - | 37 | brain | - | 21 | - | 5 | - |
| | | 38 | parotid LN | - | 3.6 | - | 12 | - |
| 31 | - | 3 | parotid LN | - | 240 | - | 240 | - |
| | | 4 | brain stem | - | 33 | - | 9.1 | - |
| | | 15 | muscle | - | 59 | - | 0.3 | - |
| 32 | - | 1 | RPLN | - | 260 | - | 246 | - |
| | | 2 | brain stem | - | 25 | - | 2.5 | - |
| | | 13 | muscle | - | 71 | - | 7.5 | - |
| 313 | - | 44 | cerebrum | - | 82 | - | 4.9 | - |
| 840 | - | 9 | RPLN | - | 410 | - | 340 | - |
| | | 17 | brain stem | - | 38 | - | 4.3 | - |
| | | 22 | muscle | - | 56 | - | 0.2 | - |
