## Appendix A, Table A2 for "Prion Seeding Activity in DNA Extractions: Implications for Laboratory Biosafety"

| | | | | MagAttract DNA | | Dneasy DNA | | Tissue |
|---|---|---|---|---|---|---|---|---|
| Animal ID | Animal's CWD Status (by ELISA or IHC) | Sample ID | Tissue | RT-QuIC Results | [DNA] (ng/uL) | RT-QuIC Results | [DNA] (ng/uL) | RT-QuIC Results |
| 287 | + | 8 | brain stem | + | 69 | + | 27 | + |
| | | 16 | muscle | - | 55 | - | 10.6 | - |
| | | 19 | parotid LN | + | 286 | - | <0.1 | + |
| | | 55 | salivary gland | + | 368 | + | 228 | + |
| | | 57 | submandibular LN | + | 45 | + | 522 | + |
| 288 | + | 39 | sciatic nerve | - | 60 | - | 0.8 | - |
| | | 40 | muscle | - | 68 | - | 13 | - |
| 289 | + | 41 | sciatic nerve | - | 58 | - | 2.2 | - |
| | | 42 | muscle | - | 112 | - | 7.1 | - |
| 306 | + | 60 | ear | + | 89 | + | 226 | + |
| | | 6 | parotid LN | + | 202 | + | 154 | + |
| | | 12 | brain stem | + | 32 | + | 5 | - |
| | | 23 | muscle | - | 36 | - | 60 | - |
| 307 | + | 48 | parotid LN | + | 266 | + | 464 | + |
| | | 49 | muscle | - | 0.7 | - | 0.7 | - |
| | | 82 | mesenteric LN | + | 0.3 | + | 2.2 | + |
| | | 83 | salivary gland | + | 16 | + | 118 | - |
| | | 84 | ear | + | 164 | + | 196 | + |
| | | 85 | bone marrow | + | 80 | - | 74 | + |
| | | 86 | popliteal LN | + | 4.8 | + | 5 | + |
| | | 87 | spleen | + | 36 | + | 40 | + |
| | | 88 | rectum | + | 32 | + | 116 | + |
| | | 89 | eye | - | 0.6 | - | 0.2 | + |
| | | 91 | inguanal LN | + | 104 | + | 400 | + |
| | | 92 | prescapular LN | + | 20 | + | 17 | + |
| | | 93 | gastrohepatic LN | + | <0.1 | + | 0.3 | + |
| | | 95 | tonsil | + | 94 | + | 14 | + |
| | | 96 | third eyelid | + | 82 | + | 466 | + |
| 310 | + | 10 | brain stem | + | 73 | + | 5 | + |
| | | 11 | RPLN | + | 322 | + | 490 | + |
| | | 14 | muscle | - | 5.7 | - | 39 | - |
| | | 99 | submandibular LN | + | 236 | + | 1160 | + |
| | | 100 | third eyelid | + | 67 | + | 125 | + |
| | | 101 | parotid LN | + | 164 | + | 744 | + |
| | | 102 | tongue | - | 60 | - | 31 | + |
| | | 103 | ear | + | 48 | + | 290 | - |
| | | 105 | salivary gland | + | 5.8 | + | 108 | + |
| 312 | + | 43 | cerebrum | - | 103 | - | 8.5 | - |
| 316 | + | 5 | RPLN | + | 742 | + | 408 | + |
| | | 18 | brain stem | - | 110 | - | 11 | - |
| | | 21 | muscle | - | 50.4 | - | 8.6 | - |
| | | 63 | parotid LN | + | 23 | + | 148 | + |

| | | | | | | | | |
|---|---|---|---|---|---|---|---|---|
| | | 64 | brain | - | 47 | + | 5.3 | + |
| | | 65 | tongue | + | 124 | + | 158 | + |
| 319 | + | 7 | brain Stem | + | 41 | + | 9.2 | + |
| | | 20 | parotid LN | + | 118 | + | 524 | + |
| | | 24 | muscle | - | 66 | - | 0.1 | - |
| | | 74 | third eyelid | + | 119 | + | 137 | + |
| | | 75 | tonsil | + | 160 | + | 240 | + |
| | | 77 | popliteal LN | + | 185 | + | 1620 | + |
| | | 78 | submandibular LN | + | 564 | + | 568 | + |
| | | 80 | brain | + | 55 | + | 0.3 | + |
| | | 81 | salivary gland | + | 96 | + | 154 | - |
