## Appendix A, Table A3 for "Prion Seeding Activity in DNA Extractions: Implications for Laboratory Biosafety"

| | | DNeasy DNA Extraction | | Tissue |
|---|---|---|---|---|
| **Sample ID** | **Experimental Status (by ELISA or IHC)** | **RT-QuIC Results** | **[DNA] (ng/uL)** | **RT-QuIC Results** |
| HB3 | control (negative) | - | 7.8 | - |
| HB6 | control (negative) | - | 7.6 | - |
| HB8 | control (negative) | - | 4.9 | - |
| HB9 | control (negative) | - | 4 | - |
| HB11 | control (negative) | - | 12 | - |
| HB13 | control (negative) | - | 4.6 | - |
| HB15 | control (negative) | - | 6.3 | - |
| HB16 | control (negative) | - | 7.4 | - |
| HB17 | control (negative) | - | 8.2 | - |
| HB4 | 80 d post inoculation | + | 5.4 | + |
| HB5 | 80 d post inoculation | + | 4.6 | + |
| HB10 | 120 d post inoculation | + | 9.9 | + |
| HB12 | 120 d post inoculation | + | 9.1 | + |
| HB14 | 120 d post inoculation | + | 6.1 | + |
| HB2 | 160 d post inoculation | + | 11 | + |
| HB7 | 160 d post inoculation | + | 8.6 | + |
| HB18 | 160 d post inoculation | + | 8.3 | + |
