## Appendix A, Table A4 for "Prion Seeding Activity in DNA Extractions: Implications for Laboratory Biosafety"

| Sample ID | DNA extraction kit | Animal CWD status (by ELISA or IHC) | DNA status | DNA concentration (ng/uL) | Dilution | Tissue RT-QuIC Result | DNA RT-QuIC Result | Tissue type |
|---|---|---|---|---|---|---|---|---|
| 82 | DNeasy | + | + | 2.2 | $10^{-1}$ | NT | + | Mesenteric lymph node |
| | | | | | $10^{-2}$ | NT | + | |
| | | | | | $10^{-3}$ | + | - | |
| | | | | | $10^{-4}$ | + | - | |
| | | | | | $10^{-5}$ | + | - | |
| | | | | | $10^{-6}$ | - | NT | |
| | | | | | $10^{-7}$ | - | NT | |
| 86 | DNeasy | + | + | 5 | $10^{-1}$ | NT | - | Popliteal lymph node |
| | | | | | $10^{-2}$ | NT | - | |
| | | | | | $10^{-3}$ | + | - | |
| | | | | | $10^{-4}$ | + | - | |
| | | | | | $10^{-5}$ | + | - | |
| | | | | | $10^{-6}$ | + | NT | |
| | | | | | $10^{-7}$ | - | NT | |
| 92 | DNeasy | + | + | 17 | $10^{-1}$ | NT | + | Prescapular lymph node |
| | | | | | $10^{-2}$ | NT | - | |
| | | | | | $10^{-3}$ | + | - | |
| | | | | | $10^{-4}$ | + | - | |
| | | | | | $10^{-5}$ | + | - | |
| | | | | | $10^{-6}$ | + | NT | |
| | | | | | $10^{-7}$ | - | NT | |
| | | | | | $10^{-8}$ | - | NT | |
| 7 | MagAttract | + | + | 41 | $10^{-1}$ | NT | + | Brain stem |
| | | | | | $10^{-2}$ | NT | - | |
| | | | | | $10^{-3}$ | + | - | |
| | | | | | $10^{-4}$ | + | - | |
| | | | | | $10^{-5}$ | + | - | |
| | | | | | $10^{-6}$ | - | NT | |
| | | | | | $10^{-7}$ | - | NT | |
| 74 | MagAttract | + | + | 119 | $10^{-1}$ | NT | + | Third eyelid |
| | | | | | $10^{-2}$ | NT | + | |
| | | | | | $10^{-3}$ | + | - | |
| | | | | | $10^{-4}$ | + | - | |
| | | | | | $10^{-5}$ | + | - | |
| | | | | | $10^{-6}$ | + | NT | |
| | | | | | $10^{-7}$ | - | NT | |
| | | | | | $10^{-8}$ | - | NT | |
| 6 | MagAttract | + | + | 202 | $10^{-1}$ | NT | + | Parotid |
| | | | | | $10^{-2}$ | NT | + | |
| | | | | | $10^{-3}$ | + | + | |
| | | | | | $10^{-4}$ | + | - | |
| | | | | | $10^{-5}$ | + | - | |
| | | | | | $10^{-6}$ | + | NT | |
| | | | | | $10^{-7}$ | - | NT | |

| | | | | | | | | |
|---|---|---|---|---|---|---|---|---|
| 99 | MagAttract | + | + | 236 | $10^{-1}$ | NT | + | Submandibular lymph node |
| | | | | | $10^{-2}$ | NT | - | |
| | | | | | $10^{-3}$ | + | + | |
| | | | | | $10^{-4}$ | + | - | |
| | | | | | $10^{-5}$ | + | - | |
| | | | | | $10^{-6}$ | + | NT | |
| | | | | | $10^{-7}$ | - | NT | |
| 20 | DNeasy | + | + | 524 | $10^{-1}$ | NT | + | Parotid |
| | | | | | $10^{-2}$ | NT | + | |
| | | | | | $10^{-3}$ | + | + | |
| | | | | | $10^{-4}$ | + | - | |
| | | | | | $10^{-5}$ | + | - | |
| | | | | | $10^{-6}$ | + | NT | |
| | | | | | $10^{-7}$ | + | NT | |
| | | | | | $10^{-8}$ | - | NT | |
| 99 | DNeasy | + | + | 1,160 | $10^{-1}$ | NT | + | Submandibular lymph node |
| | | | | | $10^{-2}$ | NT | + | |
| | | | | | $10^{-3}$ | + | + | |
| | | | | | $10^{-4}$ | + | + | |
| | | | | | $10^{-5}$ | + | - | |
| | | | | | $10^{-6}$ | + | NT | |
| | | | | | $10^{-7}$ | - | NT | |
| 43 | MagAttract | + | - | 103 | $10^{-1}$ | NT | - | Cerebrum |
| | | | | | $10^{-2}$ | NT | - | |
| | | | | | $10^{-3}$ | - | - | |
| | | | | | $10^{-4}$ | - | - | |
| | | | | | $10^{-5}$ | - | - | |
| | | | | | $10^{-6}$ | - | NT | |
| | | | | | $10^{-7}$ | - | NT | |
