## Appendix B, Table B1 for "Prion Seeding Activity in DNA Extractions: Implications for Laboratory Biosafety"

| Gradient dilution (CWD+ in CWD-) | Obex | | RPLN | |
|---|---|---|---|---|
| | DNA#1 (ng/µL) | DNA#2 (ng/µL) | DNA#1 (ng/µL) | DNA#2 (ng/µL) |
| CWD+ | 59.97 | 44.21 | 43.65 | 48.04 |
| $10^{-1}$ | 19.59 | 17.74 | 45.03 | 5.94 |
| $10^{-2}$ | 24 | 27.57 | 41.15 | 43.26 |
| $10^{-3}$ | 12.23 | 13.89 | 42.71 | 43.43 |
| $10^{-4}$ | 6.21 | 6.39 | 53.66 | 48.01 |
| $10^{-5}$ | 10.91 | 9.82 | 44.77 | 41.57 |
| CWD - | 10.71 | 11.94 | 48.23 | 43.29 |
