## Appendix B, Table B2 for "Prion Seeding Activity in DNA Extractions: Implications for Laboratory Biosafety"

| Sample ID | Tissue type | ELISA | OD value | IHC | DNA extraction KIT |
|---|---|---|---|---|---|
| 20RC0080 | Obex | POS | NA | POS | QIAcube HT |
| 20RC0081 | Obex | POS | 0.776 | LN POS/ OB NEG | QIAcube HT |
| 20RC0083 | Obex | POS | NA | POS | QIAcube HT |
| 20RC0084 | Obex | POS | NA | POS | QIAcube HT |
| 20RC0085 | Obex | POS | NA | POS | QIAcube HT |
| 20RC0088 | Obex | POS | NA | POS | QIAcube HT |
| 20RC0089 | Obex | POS | NA | POS | QIAcube HT |
| 20RC0091 | Obex | POS | NA | POS | QIAcube HT |
| 22RC0054 | Obex | NEG | 0.024 | NEG | QIAcube HT |
| 22RC2286 | Obex | NEG | 0.009 | NA | QIAcube HT |
| 22RC2360 | Obex | NEG | 0.014 | NA | QIAcube HT |
| 22RC2362 | Obex | NEG | 0.018 | NA | QIAcube HT |
| 22RC2391 | Obex | NEG | 0.018 | NA | QIAcube HT |
| 22RC2415 | Obex | NEG | 0.023 | NA | QIAcube HT |
| 22RC2422 | Obex | NEG | 0.008 | NA | QIAcube HT |
| 22RC2433 | Obex | NEG | 0.015 | NA | QIAcube HT |
| 22RC2434 | Obex | NEG | 0.014 | NA | QIAcube HT |
| 22RC2437 | Obex | NEG | 0.016 | NA | QIAcube HT |
| 22RC2588 | Obex | NEG | 0.013 | NA | QIAcube HT |
| 22RC2589 | Obex | NEG | 0.013 | NA | QIAcube HT |
| 22RC2594 | Obex | NEG | 0.013 | NA | QIAcube HT |
| 22RC2621 | Obex | NEG | 0.011 | NA | QIAcube HT |
| 22RC2648 | Obex | NEG | 0.004 | NA | QIAcube HT |
| 22RC2746 | Obex | NEG | 0.018 | NA | QIAcube HT |
| 22RC2775 | Obex | NEG | 0.031 | NA | QIAcube HT |
| 22RC2808 | Obex | NEG | 0.011 | NA | QIAcube HT |
| 22RC3019 | Obex | NEG | 0.006 | NA | QIAcube HT |
| 22RC3274 | Obex | NEG | 0.008 | NA | QIAcube HT |
| 22RC3328 | Obex | POS | NA | POS | QIAcube HT |
| 22RC3331 | Obex | POS | NA | POS | QIAcube HT |
| 22RC3335 | Obex | POS | NA | POS | QIAcube HT |
| 22RC3340 | Obex | POS | NA | LN POS/ OB NEG | QIAcube HT |
| 22RC3346 | Obex | POS | NA | POS | QIAcube HT |
| 22RC3348 | Obex | POS | 0.053 | POS | QIAcube HT |
| 22RC3351 | Obex | POS | NA | POS | QIAcube HT |
| 22RC3355 | Obex | POS | NA | POS | QIAcube HT |
| 22RC3362 | Obex | POS | NA | LN POS/ OB NEG | DNeasy |
| 22RC3580 | Obex | POS | NA | POS | QIAcube HT |

| | | | | | |
|---|---|---|---|---|---|
| 22RC3586 | Obex | POS | NA | POS | QIAcube HT |
| 22RC3590 | Obex | POS | NA | POS | QIAcube HT |
| 22RC3594 | Obex | POS | NA | POS | QIAcube HT |
| 22RC3600 | Obex | POS | NA | POS | QIAcube HT |
| 22RC3601 | Obex | POS | NA | POS | QIAcube HT |
| 22RC3604 | Obex | POS | NA | POS | QIAcube HT |
| 22RC3605 | Obex | POS | NA | LN POS/ OB NEG | QIAcube HT |
| 22RC3611 | Obex | POS | NA | POS | QIAcube HT |
| 22RC3612 | Obex | POS | NA | POS | QIAcube HT |
| 22RC3622 | Obex | POS | NA | POS | QIAcube HT |
| 24RC0005 | RPLN | NEG | 0.02 | NA | QIAcube HT |
| 24RC0006 | RPLN | NEG | 0.013 | NA | QIAcube HT |
| 24RC0007 | RPLN | NEG | 0.015 | NA | QIAcube HT |
| 24RC0008 | RPLN | NEG | 0.013 | NA | QIAcube HT |
| 24RC0009 | RPLN | NEG | 0.014 | NA | QIAcube HT |
| 24RC0010 | RPLN | NEG | 0.012 | NA | QIAcube HT |
| 24RC0011 | RPLN | NEG | 0.015 | NA | QIAcube HT |
| 24RC0012 | RPLN | NEG | 0.017 | NA | QIAcube HT |
| 24RC0013 | RPLN | NEG | 0.022 | NA | QIAcube HT |
| 24RC0014 | RPLN | NEG | 0.014 | NA | QIAcube HT |
| 24RC0015 | RPLN | NEG | 0.011 | NA | QIAcube HT |
| 24RC0016 | RPLN | NEG | 0.005 | NA | QIAcube HT |
| 24RC0200 | RPLN | POS | NA | POS | QIAcube HT |
| 24RC0201 | RPLN | POS | NA | POS | QIAcube HT |
| 24RC0202 | RPLN | POS | NA | POS | QIAcube HT |
| 24RC0204 | RPLN | POS | NA | POS | QIAcube HT |
| 24RC0205 | RPLN | POS | NA | POS | QIAcube HT |
| 24RC0206 | RPLN | POS | NA | POS | QIAcube HT |
| 24RC0207 | RPLN | POS | NA | POS | QIAcube HT |
| 24RC0208 | RPLN | POS | NA | POS | QIAcube HT |
| 24RC0210 | RPLN | POS | NA | POS | QIAcube HT |
| 24RC0211 | RPLN | POS | NA | POS | QIAcube HT |
| 24RC0212 | RPLN | POS | NA | POS | QIAcube HT |
| 24RC0213 | RPLN | POS | NA | POS | QIAcube HT |
| 24RC0214 | RPLN | POS | NA | POS | QIAcube HT |
| 24RC0215 | RPLN | POS | NA | POS | QIAcube HT |
| 24RC0216 | RPLN | POS | NA | POS | QIAcube HT |
| 24RC0227 | RPLN | POS | 3.5 | POS | DNeasy |
| 24RC0278 | RPLN | POS | NA | POS | QIAcube HT |
| 24RC0279 | RPLN | POS | NA | POS | QIAcube HT |
| 24RC0280 | RPLN | POS | NA | POS | QIAcube HT |
| 24RC0281 | RPLN | POS | NA | POS | QIAcube HT |

| | | | | | |
|---|---|---|---|---|---|
| 24RC0283 | RPLN | POS | NA | POS | QIAcube HT |
| 24RC0289 | RPLN | POS | NA | POS | QIAcube HT |
| 24RC0339 | RPLN | POS | NA | POS | QIAcube HT |
| 24RC0342 | RPLN | POS | NA | POS | QIAcube HT |
| 24RC0358 | RPLN | POS | NA | POS | DNeasy |
| 24RC0017 | RPLN | NEG | NA | NA | QIAcube HT |
| 24RC0019 | RPLN | NEG | NA | NA | QIAcube HT |
| 24RC0020 | RPLN | NEG | NA | NA | QIAcube HT |
