## Appendix B, Table B3 for "Prion Seeding Activity in DNA Extractions: Implications for Laboratory Biosafety"

| Sample name | # RT-QuIC replicates | # POS replicates (CT < 30h) | Cycle Threshold (CT) for individual replicate | | | | | | | |
|---|---|---|---|---|---|---|---|---|---|---|
| 20RC0080 | 4 | 4/4 | 11.61 | 13.24 | 11.45 | 10.25 | | | | |
| 20RC0081 | 8 | 0/8 | 65 | 65 | 65 | 55.07 | 56.12 | 65 | 46.28 | 65 |
| 20RC0083 | 8 | 4/8 | 8.91 | 65 | 11.65 | 11.61 | 43.39 | 69.4 | 65 | 11.03 |
| 20RC0084 | 8 | 7/8 | 65 | 8.24 | 9.24 | 14.44 | 7.61 | 6.76 | 5.21 | 6.09 |
| 20RC0085 | 4 | 4/4 | 8.35 | 7.35 | 13.25 | 9.7 | | | | |
| 20RC0088 | 8 | 1/8 | 31.25 | 47.61 | 23.14 | 65 | 32.4 | 77.92 | 65 | 65 |
| 20RC0089 | 8 | 2/8 | 43.42 | 8.23 | 11.89 | 57.64 | 65 | 48.6 | 65 | 65 |
| 20RC0091 | 8 | 1/8 | 65 | 65 | 16.15 | 30.26 | 65 | 64.28 | 75.75 | 65 |
| 22RC0054 | 4 | 0/4 | 65 | 65 | 65 | 65 | | | | |
| 22RC2286 | 4 | 0/4 | 65 | 65 | 65 | 65 | | | | |
| 22RC2360 | 4 | 0/4 | 65 | 65 | 65 | 65 | | | | |
| 22RC2362 | 4 | 0/4 | 65 | 65 | 65 | 65 | | | | |
| 22RC2391 | 4 | 0/4 | 65 | 65 | 65 | 65 | | | | |
| 22RC2415 | 4 | 0/4 | 65 | 65 | 65 | 65 | | | | |
| 22RC2422 | 4 | 0/4 | 65 | 65 | 65 | 65 | | | | |
| 22RC2433 | 4 | 0/4 | 65 | 65 | 65 | 65 | | | | |
| 22RC2434 | 4 | 0/4 | 65 | 65 | 65 | 65 | | | | |
| 22RC2437 | 4 | 0/4 | 65 | 65 | 65 | 40.22 | | | | |
| 22RC2588 | 4 | 0/4 | 65 | 65 | 65 | 65 | | | | |
| 22RC2589 | 8 | 0/8 | 65 | 65 | 65 | 40.6 | 45.6 | 41.61 | 69.67 | 65 |
| 22RC2594 | 8 | 4/8 | 65 | 42.45 | 22.35 | 65 | 21.69 | 12.45 | 13.49 | 65 |
| 22RC2621 | 8 | 2/8 | 65 | 5.98 | 31.29 | 65.56 | 65 | 25.38 | 30.65 | 65 |
| 22RC2648 | 8 | 0/8 | 65 | 62.45 | 53.35 | 68.3 | 65 | 65 | 65 | 65 |
| 22RC2746 | 8 | 1/8 | 65 | 65 | 65 | 38.26 | 23.49 | 41.42 | 65 | 65 |
| 22RC2775 | 4 | 0/4 | 65 | 65 | 65 | 65 | | | | |
| 22RC2808 | 8 | 2/8 | 19.21 | 65 | 27.05 | 65 | 78.55 | 31.13 | 65 | 65 |
| 22RC3019 | 8 | 2/8 | 34.62 | 65 | 65 | 34.45 | 29.36 | 28.51 | 36.38 | 41.61 |
| 22RC3274 | 8 | 2/8 | 65 | 65 | 43.18 | 65 | 19.25 | 26.69 | 31.27 | 35.55 |
| 22RC3328 | 8 | 2/8 | 65 | 65 | 42.65 | 31.7 | 65 | 9.72 | 41.49 | 9.69 |
| 22RC3331 | 8 | 6/8 | 24.39 | 21.65 | 52.42 | 65 | 11.46 | 18.29 | 11.97 | 14.24 |
| 22RC3335 | 8 | 7/8 | 16.72 | 21.73 | 65 | 16.63 | 13.31 | 10.15 | 13.09 | 10.23 |
| 22RC3340 | 8 | 4/8 | 68.47 | 35.27 | 65 | 17.11 | 19.26 | 49.47 | 24.3 | 19.06 |
| 22RC3346 | 8 | 5/8 | 23.47 | 28.45 | 65 | 43.17 | 3.72 | 6.19 | 37.7 | 6.18 |
| 22RC3348 | 4 | 4/4 | 7.68 | 18.01 | 18.59 | 8.83 | | | | |
| 22RC3351 | 4 | 4/4 | 9.2 | 17.27 | 12.4 | 8.95 | | | | |
| 22RC3355 | 4 | 4/4 | 6.83 | 13.76 | 14.71 | 10.44 | | | | |
| 22RC3362 | 4 | 4/4 | 8.74 | 16.25 | 11.52 | 12.55 | | | | |
| 22RC3580 | 8 | 2/8 | 42.07 | 65 | 65 | 40.37 | 20.26 | 8.95 | 65 | 65 |
| 22RC3586 | 8 | 2/8 | 41.42 | 48.74 | 65 | 65 | 19.26 | 9.54 | 66.32 | 65 |
| 22RC3590 | 4 | 0/4 | 65 | 65 | 65 | 65 | | | | |
| 22RC3594 | 4 | 0/4 | 65 | 65 | 65 | 65 | | | | |
| 22RC3600 | 8 | 6/8 | 9.32 | 65 | 19.31 | 18.83 | 9.96 | 8.14 | 65 | 16.54 |
| 22RC3601 | 4 | 4/4 | 9.29 | 14.34 | 17.28 | 18.66 | | | | |
| 22RC3604 | 4 | 4/4 | 9.74 | 15.62 | 13.49 | 14.52 | | | | |
| 22RC3605 | 8 | 6/8 | 31.69 | 14.24 | 12.45 | 9.24 | 65 | 21.3 | 19.59 | 22.43 |
| 22RC3611 | 8 | 6/8 | 65 | 33.18 | 17.28 | 19.9 | 14.25 | 8.9 | 7.33 | 25.47 |
| 22RC3612 | 8 | 4/8 | 65 | 16.25 | 27.76 | 31.27 | 37.51 | 12.24 | 65 | 8.07 |
| 22RC3622 | 8 | 6/8 | 65 | 65 | 13.44 | 14.55 | 9.08 | 5.81 | 14.08 | 8.72 |
| 24RC0005 | 4 | 0/4 | 65 | 65 | 65 | 65 | | | | |

| | | | | | | | | | | |
|---|---|---|---|---|---|---|---|---|---|---|
| 24RC0006 | 8 | 1/8 | 65 | 65 | 27.64 | 65 | 65 | 78.43 | 65 | 65 |
| 24RC0007 | 8 | 0/8 | 65 | 38.02 | 65 | 65 | 65 | 65 | 65 | 65 |
| 24RC0008 | 4 | 0/4 | 65 | 65 | 65 | 65 | | | | |
| 24RC0009 | 4 | 0/4 | 65 | 65 | 65 | 65 | | | | |
| 24RC0010 | 4 | 0/4 | 65 | 65 | 65 | 65 | | | | |
| 24RC0011 | 8 | 2/8 | 65 | 22.32 | 14.71 | 32.55 | 65 | 65 | 65 | 65 |
| 24RC0012 | 8 | 3/8 | 22.32 | 29.54 | 65 | 65 | 65 | 31.5 | 9.36 | 42.26 |
| 24RC0013 | 8 | 3/8 | 26.29 | 29.35 | 36.19 | 20.25 | 65 | 65 | 65 | 65 |
| 24RC0014 | 8 | 2/8 | 65 | 65 | 35.25 | 65 | 29.43 | 75.75 | 20.27 | 41.69 |
| 24RC0015 | 4 | 0/4 | 65 | 65 | 65 | 65 | | | | |
| 24RC0016 | 8 | 4/8 | 24.57 | 65 | 19.31 | 38.33 | 23.21 | 49.47 | 26.51 | 48.34 |
| 24RC0200 | 4 | 4/4 | 9.22 | 16.42 | 13.44 | 15.15 | | | | |
| 24RC0201 | 4 | 4/4 | 13.05 | 17.58 | 20.25 | 23.08 | | | | |
| 24RC0202 | 8 | 2/8 | 17.9 | 38.69 | 15.74 | 34.55 | 65 | 65 | 47.41 | 65 |
| 24RC0204 | 8 | 4/8 | 24.28 | 13.82 | 16.26 | 41.67 | 16.14 | 65 | 65 | 65 |
| 24RC0205 | 8 | 5/8 | 14.67 | 14.27 | 27.03 | 41.65 | 17.31 | 17.45 | 65 | 65 |
| 24RC0206 | 8 | 4/8 | 43.49 | 39.46 | 21.17 | 28.76 | 18.26 | 65 | 65 | 22.31 |
| 24RC0207 | 8 | 3/8 | 31.45 | 32.7 | 16.87 | 65 | 22.03 | 20.51 | 45.75 | 65 |
| 24RC0208 | 4 | 0/4 | 65 | 65 | 65 | 65 | | | | |
| 24RC0210 | 4 | 4/4 | 8.23 | 13.23 | 11.2 | 7.13 | | | | |
| 24RC0211 | 4 | 4/4 | 10.23 | 6.75 | 8.59 | 7.13 | | | | |
| 24RC0212 | 8 | 4/8 | 9.15 | 28.37 | 20.37 | 12.57 | 39.42 | 65 | 65 | 65 |
| 24RC0213 | 4 | 4/4 | 8.72 | 6.58 | 7.31 | 8.74 | | | | |
| 24RC0214 | 8 | 4/8 | 19.28 | 16.39 | 65 | 65 | 25.26 | 65 | 38.72 | 15.2 |
| 24RC0215 | 4 | 4/4 | 13.51 | 14.24 | 12.22 | 10.48 | | | | |
| 24RC0216 | 8 | 1/8 | 65 | 65 | 65 | 20.4 | 46.62 | 57.82 | 65 | 68.3 |
| 24RC0227 | 8 | 5/8 | 9.65 | 65 | 15.17 | 65 | 3.36 | 7.22 | 11.19 | 65 |
| 24RC0278 | 8 | 5/8 | 13.57 | 65 | 19.52 | 44.44 | 14.74 | 45.25 | 22.85 | 12.31 |
| 24RC0279 | 8 | 5/8 | 14.21 | 23.27 | 10.25 | 65 | 37.49 | 12.63 | 38.59 | 23.28 |
| 24RC0280 | 8 | 5/8 | 13.33 | 11.17 | 29.35 | 19.43 | 65 | 44.28 | 10.91 | 65 |
| 24RC0281 | 4 | 4/8 | 14.24 | 17.87 | 12.31 | 12.27 | | | | |
| 24RC0283 | 8 | 6/8 | 16.47 | 14.24 | 29.27 | 65 | 10.28 | 21.61 | 14.59 | 67.29 |
| 24RC0289 | 8 | 8/8 | 10.19 | 13.23 | 21.54 | 25.4 | 14.08 | 10.21 | 28.66 | 19.1 |
| 24RC0339 | 8 | 7/8 | 23.49 | 14.33 | 33.4 | 25.38 | 12.48 | 17.56 | 19.27 | 18.28 |
| 24RC0342 | 4 | 4/4 | 6.72 | 13.22 | 17.7 | 7.13 | | | | |
| 24RC0358 | 4 | 0/4 | 65 | 65 | 65 | 65 | | | | |
| 24RC0017 | 8 | 0/8 | 65 | 65 | 65 | 65 | 65 | 65 | 65 | 65 |
| 24RC0019 | 8 | 0/8 | 65 | 65 | 65 | 65 | 65 | 65 | 65 | 65 |
| 24RC0020 | 8 | 2/8 | 65 | 28.5 | 65 | 29.27 | 65 | 65 | 65 | 65 |
