## Appendices A and B text for "Prion Seeding Activity in DNA Extractions: Implications for Laboratory Biosafety"

### Appendix A

<u>MNPRO Supplemental Data</u>

**Table A1.** MNPRO RT-QuIC results for tissue and both types of DNA eluates in all CWD-negative WTD samples used in these experiments.

**Table A2.** MNPRO RT-QuIC results for tissue and both types of DNA eluates in all MNPRO CWD-positive WTD samples used in these experiments.

**Table A3.** MNPRO RT-QuIC results for tissue and DNA eluates for all MNPRO Syrian hamster (*Mesocricetus auratus*) samples used in these experiments.

**Table A4.** MNPRO RT-QuIC results for endpoint titration experiments. NT = not tested. Sample IDs here correspond with the Sample IDs shown in Table A2.

### Appendix B

<u>CFIA Supplemental Data</u>

**Table B1. Concentration of DNA extracted from control tissue homogenates[1]**
[1]Tissue homogenates were mixtures of CWD+ and CWD- tissue homogenate in various ratios, each containing 0.01% (w/v) of tissue specimens. DNA was extracted from the tissue homogenates using the DNeasy Blood & Tissue kit (Qiagen) and quantified using the Quant-iT kit (Invitrogen).

**Table B2**. CFIA sample information

**Table B3.** RT-QuIC data for DNA samples

**Figure B1**. Correlation between the concentration and cycle threshold in DNA samples
