## Supplementary figures and images for "Prion Seeding Activity in DNA Extractions: Implications for Laboratory Biosafety"

### Appendix B, Figure B1

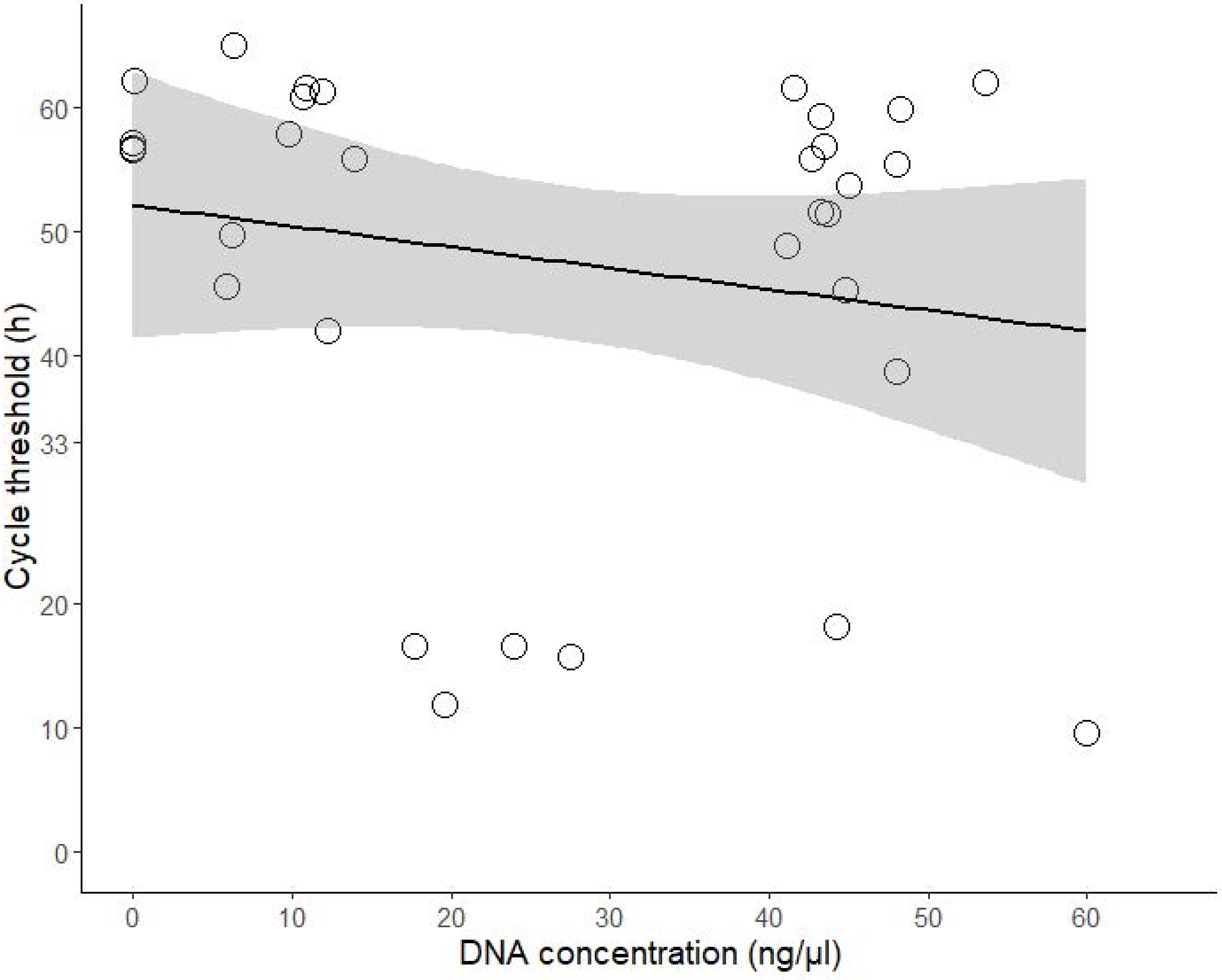
